# The Spatial Architecture of Human Subgingival Biofilms

**DOI:** 10.64898/2026.09.19.752862

**Authors:** Ananya Padmakumar, Paul J McMillan, HoJin Chang, Luan Ngo, Harikrishnan Parappalliyalil, Shafira Bianca Susilo, Jeremy D. Silver, Stuart G. Dashper, Eric C. Reynolds, Paul D. Veith, Debnath Ghosal

## Abstract

Human microbiome studies have transformed understanding of disease-associated microbial communities, yet most approaches collapse polymicrobial biofilms into compositional profiles and lose the spatial relationships through which microorganisms interact *in vivo*. Periodontitis provides a clinically important model to investigate microbial spatial organisation because the disease is associated with deep, anaerobic periodontal pockets colonized by dense polymicrobial biofilms. Here, using multiplex fluorescence *in situ* hybridization with spectral and confocal microscopy of subgingival plaque across 174 sites from 36 individuals with periodontitis, we define taxon-resolved spatial architectures of the human subgingival plaque microbiome. We show that subgingival plaque is organized into recurrent multicellular assemblages of microbial consortia rather than unstructured layers of biomass, including assemblages defined as test-tube-brushes, corncobs and additional polymicrobial architectures with distinct taxonomic patterning. Test-tube brushes were enriched in severe periodontal niches, and their detection and burden increased with pocket depth, with distinct patterns observed in smoking-associated disease sites. In a longitudinal treatment subset, sites containing test-tube brushes before therapy lacked detectable test-tube brushes after treatment and instead showed corncobs concomitant with reduced pocket depth and resolution of disease. In addition to canonical oral biofilm structures, we identify previously undescribed polymicrobial assemblages whose occurrence is patterned by pocket depth and anatomical tooth site. These findings show that disease-associated human biofilms are organized into reproducible spatial states and suggest that microbial architecture provides an additional axis for interpreting microbiome-associated disease.

**Graphical Abstract:** 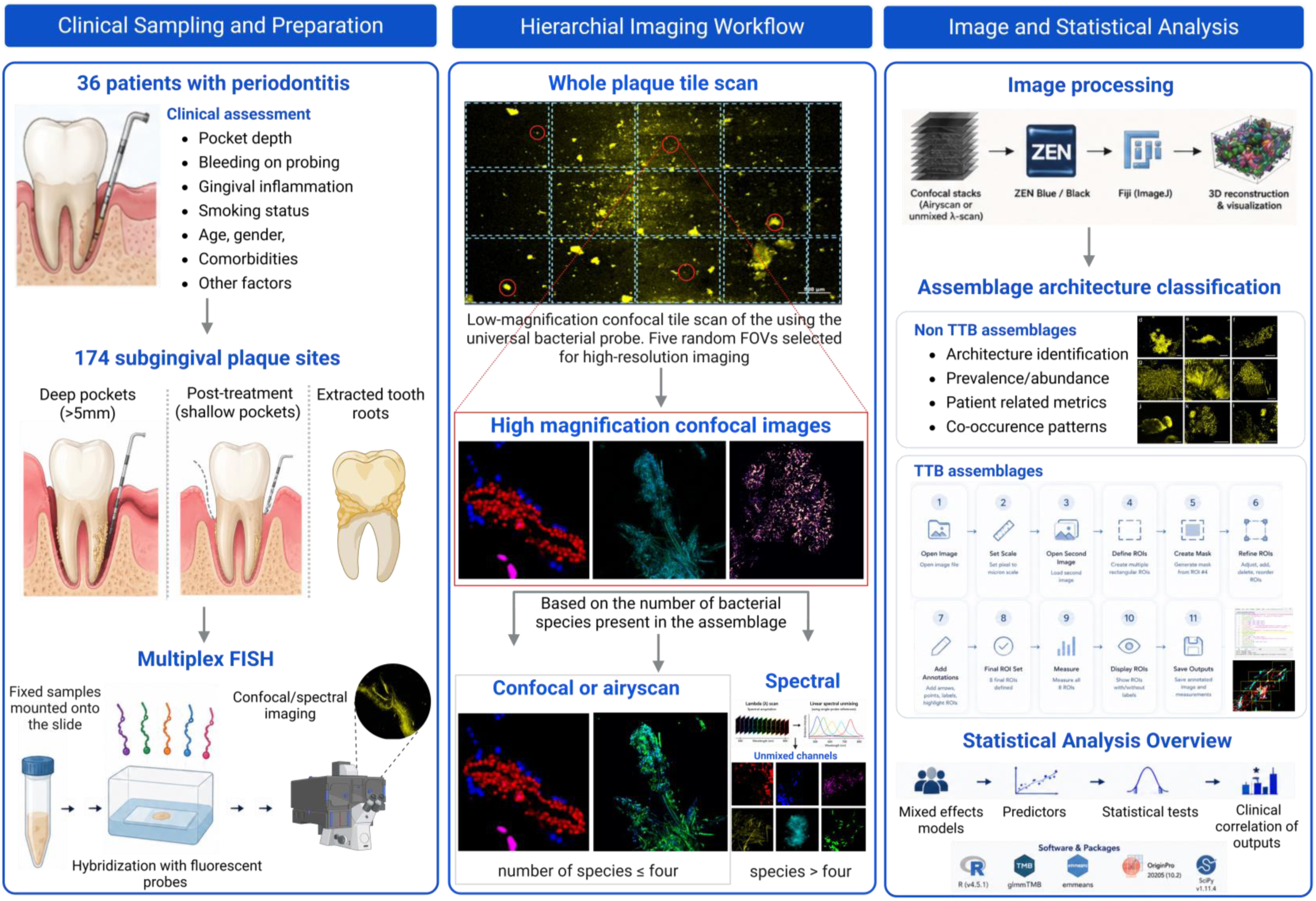

## Introduction

Microbiome studies have transformed understanding of human disease by identifying microbial taxa and community configurations associated with health and pathology. However, in polymicrobial biofilm-associated diseases, community composition alone may not capture the spatial organisation through which microorganisms interact *in vivo*. Spatial organisation is a fundamental determinant of biofilm function, shaping microbial metabolism, interspecies interactions, nutrient exchange, local microenvironments and exposure to host-derived stress^1^. In disease-associated biofilms, this organisation may influence community function, clinical progression and treatment response. Yet for many human-associated polymicrobial biofilms, the taxon-resolved architectures that form in *vivo* and their relationship to disease severity remain poorly defined.

Periodontitis provides a clinically important model to investigate this issue of polymicrobial spatial organisation. It is among the most prevalent chronic inflammatory and destructive diseases of humans, causing tooth loss, impaired oral function and reduced quality of life^2^. At the centre of periodontitis is subgingival plaque, a dense polymicrobial biofilm that develops below the gingival margin within nutrient-rich, inflamed and increasingly anaerobic periodontal pockets^3–6^. Sequencing studies have extensively defined the taxonomic and functional composition of subgingival plaque, revealing reproducible community shifts associated with disease severity, inflammation and treatment^7–10^. However, these approaches largely collapse biofilms into bulk molecular profiles and obscure a fundamental feature of microbial communities: their three-dimensional organisation.

Spatial organisation is central to oral biofilm biology. In supragingival plaque, fluorescence *in situ* hybridisation (FISH) and spectral imaging have revealed conserved, taxonomically ordered architectures such as corncobs and hedgehogs, establishing that dental plaque is not an amorphous microbial mass, but a structured consortium composed of reproducible multicellular assemblages^11–13^. In contrast, the architecture of subgingival plaque (SubP), particularly at the base of deep periodontal pockets associated with severe periodontitis, remains poorly resolved. Earlier ultrastructural studies^14^ provided important morphological views of tooth-attached subgingival deposits but lacked taxon-resolved spatial information. To date, one of the few FISH-based taxon-resolved studies of natural subgingival biofilm architecture examined teeth from four subjects but did not systematically relate spatial structures to probing depth, smoking exposure, disease severity or treatment outcome ^15^.

This is a critical gap because deep periodontal pockets represent ecological niches directly linked to tissue destruction, anaerobiosis, inflammatory exudate and clinical progression^8^. Although structural assemblages such as corncobs and test-tube-brush structures have been observed in oral biofilms, their recurrence, cellular composition and clinical significance in subgingival plaque remain undefined. A structure-resolved framework is therefore required to determine whether biofilm architecture represents an additional axis of periodontal disease biology, complementary to microbial composition and host inflammatory state.

Here, we used multiplex FISH with spectral and confocal microscopy to analyse SubP from 174 sites across 36 individuals with periodontitis. We show that the human SubP microbiome is organised into recurrent taxon-resolved multicellular architectures rather than spatially unstructured biomass. Among these, test-tube-brush structures formed ordered polymicrobial assemblages with filamentous cores and radially arranged bacterial cells, were enriched in severe periodontal niches, and showed increased detection and burden with pocket depth. In a longitudinal treatment subset, sites containing test-tube brushes before therapy lacked detectable test-tube brushes after treatment and instead showed corncobs concomitant with reduced pocket depth and resolution of disease. In addition to canonical oral biofilm structures, we identify previously undescribed polymicrobial assemblages whose occurrence is patterned by pocket depth and anatomical tooth site. Together, these findings identify spatial architecture as an additional disease-state axis in the human subgingival microbiome.

## Results

### Clinical profiling captures heterogeneous periodontal disease niches

We analyzed 174 subgingival plaque (SubP) samples from 36 periodontitis patients to define the diversity and clinical patterning of taxon-resolved microbial assemblages within a clinically stratified cohort (Supplementary Table 1). We have used the FDI World Dental Federation Notation (ISO 3950) where 13MD (9mm) refers to a 9 mm periodontal pocket mesiodistal to the canine (tooth 3) of the upper right quadrant (1) (Extended Data Fig. 1a). Clinical descriptors are reported to contextualize the imaging dataset (Fig. 1a-c). To identify the most informative clinical metric for stratifying sites in our imaging dataset, we quantified maximum pocket depth (PD) and bleeding-on-probing (BoP) percentage for each patient (Fig. 1a, Extended Data Fig. 1b). Spearman’s correlation revealed a weak, non-significant association between maximum PD and BoP percentage at the patient level (ρ = 0.23, p = 0.17; n = 36), indicating that bleeding did not consistently increase with deeper pockets in this cohort, consistent with subsequent site-level mixed-effects modelling. Given that PD provides a continuous site-resolved measure of structural disease burden, maximum PD (defined as the deepest site recorded per patient) was selected as the primary severity metric for stratifying the imaging dataset.

**Fig. 1.**
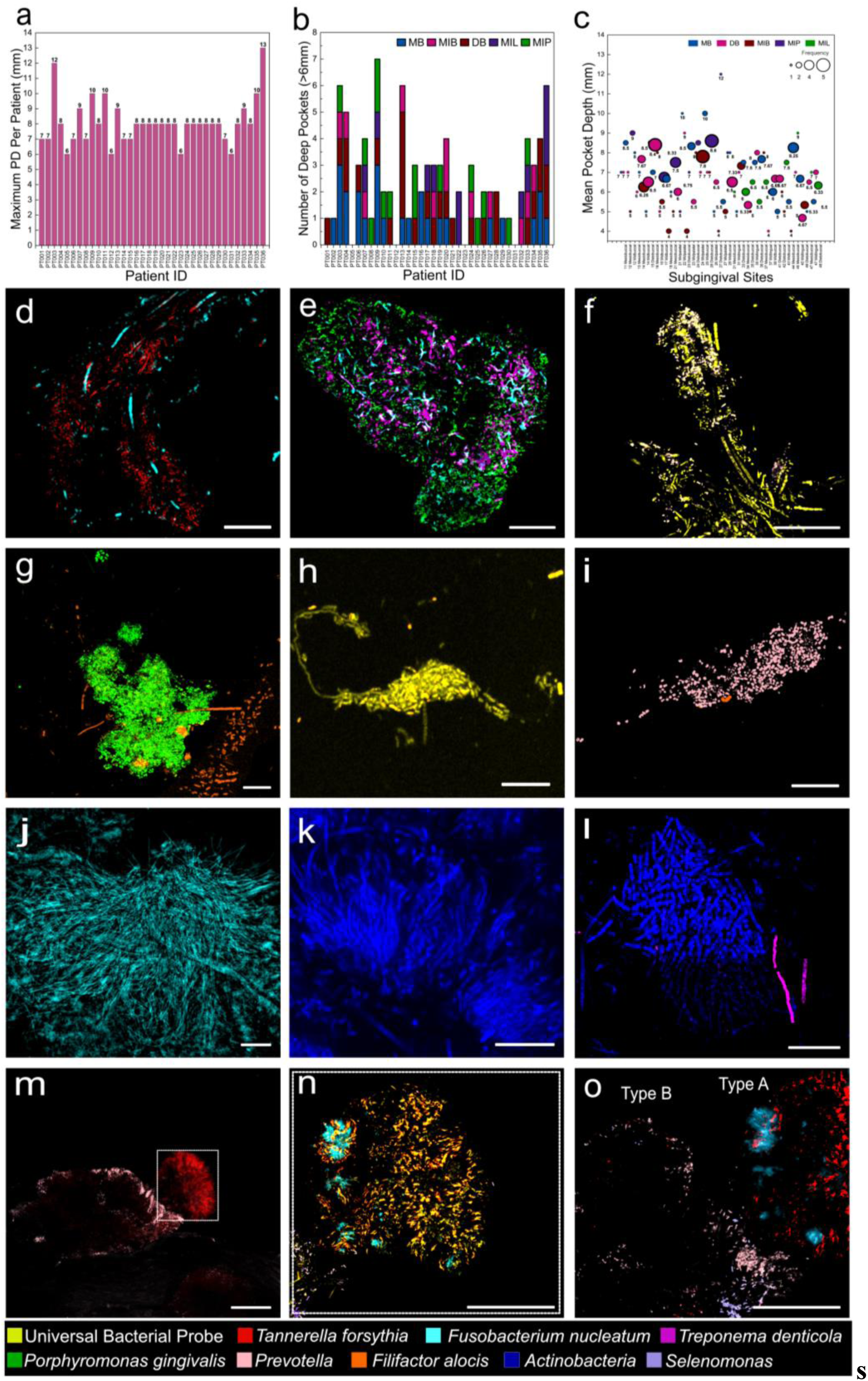
a-c) Clinical profiling and anatomical distribution of periodontal pocket depth across the imaging cohort. b-o) Distinct spatial assemblages of key periodontal taxa revealed by multiplexed FISH and confocal imaging. **a)** Maximum periodontal pocket depth (mm) recorded per patient across the 36-patient periodontitis cohort. Each bar represents one patient, with the value above the bar indicating the patient-level maximum pocket depth used to stratify clinical disease severity in the imaging dataset. b) Number of sampled periodontal sites with pocket depth >6 mm for each patient, stratified by tooth site: mesiobuccal (MB), midbuccal (MIB), distobuccal (DB), midlingual (MIL) and midpalatal (MIP). Stacked bars show the contribution of each tooth site to the total number of deep periodontal sites sampled for each patient. Stacked bars show the contribution of each tooth site to the total number of deep sampled sites for each patient. c) Site-level distribution of mean pocket depth across sampled subgingival sites. Each bubble represents a sampled site and is coloured by anatomical tooth surface. Bubble size indicates the frequency of sites with the corresponding pocket depth value. The plot shows substantial heterogeneity in pocket depth across sites, with deeper pockets more frequently observed at lingual and mesial surfaces. d) TTB brush structure e) *Pg-Td-Fn* cluster f) Unique corncob-like structure from a smoker with diabetes, featuring filamentous *Fn* at the core and *Prevotella* with other eubacterial members at the periphery. g) *Achillea*-like clusters formed by *Pg* (dense central core) and *Fa* (peripheral distribution). h) Dragon-like assemblage primarily made of *Fa*. i) *Prevotella* and *Fa* dense microcolony. j) Filamentous *Fn* biostreams. k) *Echidna*-like assemblage composed of Actinobacteria, characterized by radial filamentous extensions. l)*Act*-cluster, formed by Actinobacteria. Panels correspond to maximum-intensity projections from CLSM images of SubP stained with taxon-specific FISH probes. Most assemblages were observed across multiple patients and sites, with several consistent pairings (e.g., *Tf*+*Fn*, *Pg*+*Fa*) and rarer assemblages (dragons) linked to specific taxa or patient profiles. m) Low-magnification view showing an amorphous orb. n) Low-magnification view showing an amorphous orb. o) The outer shell comprised taxonomically defined species distinguishing Type A and Type B ‘amorphous orbs. Type A, dominated by *Tf* and *Fn*; and Type B, enriched in *Prevotella* and *Selenomonas*. Extended Data Fig. 3. shows grayscale reference images for multiplexed FISH channels. Supplementary Fig. S1 shows additional probe sets and morphological variants of these assemblages. Scale bars: 10 μm.

When stratified by smoking status, smokers exhibited significantly greater pocket depths while showing similar bleeding-on-probing frequencies to non-smokers (Extended Data Fig. 2a, b). Mixed-effects linear regression showed smoking status was significantly associated with greater pocket depth (+2.53 mm, 95% CI 1.60–3.47, p < 0.0001), while mixed-effects logistic regression showed no significant association between smoking and BoP (p = 0.37). Several smokers under 50 years of age also displayed average PDs exceeding the Stage III/IV clinical disease threshold of 6 mm, indicating substantial disease burden in younger smokers in this cohort (Extended Data Fig. 2c). Consistent with these observations, mixed-effects modelling identified smoking, but not age, as a significant predictor of pocket depth.

We next examined the anatomical distribution of disease across the periodontal tooth sites represented in the imaging cohort by assessing the distribution of deep periodontal pockets (>6 mm) across patients (Fig. 1b). Across the 174 sampled periodontal sites, the imaging cohort comprised mesiobuccal (MB), distobuccal (DB), midbuccal (MIB), midlingual (MIL) and midpalatal (MIP) SubP samples. A site-level bubble plot further revealed substantial spatial heterogeneity across sampled subgingival sites, with most sites clustering between 5–8 mm and deeper pockets more frequently observed at lingual and mesial surfaces (Fig. 1c). Consistent with these site-level patterns, mixed-effects linear regression showed that anatomical tooth surface was significantly associated with pocket depth (χ2=17.23, df = 4, p=0.0017). Midlingual sites had significantly greater estimated pocket depths than distobuccal sites by 0.82 mm (95% CI, 0.04–1.60 mm; adjusted p=0.032) and midbuccal sites by 1.07 mm (95% CI, 0.24–1.90 mm; adjusted p=0.004; Supplementary Fig. S1). No other pairwise comparisons were significant after correction for multiple testing.

### Specific taxa exhibit defined spatial arrangements within subgingival assemblages

High-resolution confocal imaging (13-26 pixels/micron) with a universal bacterial FISH probe revealed a diverse spectrum of spatially distinct microbial assemblages across patients and sampling sites (Supplementary Fig S2a–l; Supplementary Table 2). These assemblages, defined here as microscale communities composed of one or more bacterial taxa, varied in geometry, composition, and prevalence. We have used the FDI World Dental Federation Notation (ISO 3950) where 13 Distal (9mm) refers to a 9 mm periodontal pocket distal to the canine (tooth 3) of the upper right quadrant (1) (Supplementary Table 1; Extended Data Fig. 1a).

The most conspicuous and recurrent assemblages were test-tube brushes (TTBs) (Fig. 1d; Extended Data Fig. 3a–c), composed of long filamentous cores surrounded by peripherally arranged shorter bacteria oriented orthogonally. These were frequently observed in deeper pockets (8–12 mm) in smokers and 10-13 mm non-smokers and often included conserved co-occurring taxa such as *Tannerella forsythia (Tf) and Fusobacteria nucleatum* (*Fn)*.

Distinct *Pg–Td–Fn* clusters were identified as mixed, multispecies assemblages in which *Porphyromonas gingivalis* (*Pg*), *Treponema denticola* (*Td*) and *Fusobacterium nucleatum* (*Fn*) co-localised within the same structure (Fig. 1e; Extended Data Fig. 3d–e; Supplementary Fig. S3a–c). These clusters were observed at tooth sites 13DB, 12DB, 15MIL, 17MIL and 16DB, spanning non-smokers, light smokers and heavy smokers, indicating that this spatial association was reproducible across multiple clinical contexts rather than being restricted to a single patient or site.

A rare corncob-like assemblage was observed at tooth site 23MB in a 10 mm periodontal pocket of a heavy smoker with diabetes (Fig. 1f). This structure contained a long filamentous core of unresolved taxonomic identity, with *Prevotella* and other eubacterial members arranged around the periphery.

Previously undescribed *Achillea*-like floral clusters were characterised by a dense *Pg* core surrounded by peripheral *Filifactor alocis* (*Fa*) cells (Fig. 1g; Extended Data Fig. 3j–l; Supplementary Fig. S3d–f). These assemblages were identified at tooth sites 26DB, 17DB, 11MB, 21MB, 26MIB, 14DB and 35DB, encompassing 5–9 mm periodontal pockets in predominantly non-smokers, with one 5 mm site from a light smoker.

A dragon-like assemblage containing *Fa* and other taxa was also observed at tooth site 26MB in a 12 mm periodontal pocket from a heavy smoker with diabetes (Fig. 1h; Extended Data Fig. 3m–o), suggesting that some architectures may be patient- or context-specific.

Compact *Prevotella* microcolonies were also detected, occasionally associated with sparse *Fa* cells (Fig. 1i; Supplementary Fig. S3g–i). These structures were observed at tooth sites 45MIL, 24MIL and 36DB, representing deep periodontal pockets in both light and heavy smokers.

Bio-streams were primarily composed of *Fn* (Fig. 1j) but were also formed by members of the orders Lactobacillales and Bacillales (Supplementary Fig. S3j), Synergistetes cluster A (Supplementary Fig. S3k), and, less frequently, Desulfobulbaceae (Supplementary Fig. S3l). These filamentous stream-like assemblages were observed at tooth sites 14DB, 16DB, 45MIL and 26MIL, spanning 6–12 mm periodontal pockets in non-smokers, light smokers and heavy smokers.

Unique *Echidna*-like assemblages consisted of Actinobacteria with radial filamentous extensions (Fig. 1k; Supplementary Fig. S2h, S3m–o). These structures were identified at tooth sites 26MB, 27DB, 16MB and 26MIL, representing 7–9 mm periodontal pockets in light smokers.

Actinobacteria-rich clusters (Act-clusters) were also observed (Fig. 1l; Extended Data Fig. 3p–r; Supplementary Fig. S3p–r), displaying variation from coccoid to elongated cellular morphologies at tooth sites 26MIL and 21MIB, corresponding to 7–9 mm periodontal pockets in non-smokers.

While conserved assemblages such as TTBs and corncobs were widespread, rarer forms, including dragons and unique corncobs (formed by *Prevotella* and other species), were often confined to specific sites, depths, or clinical profiles (e.g., heavy smoker, comorbid diabetes). These patterns highlight that SubP hosts spatially organized, taxon-specific assemblages with both reproducibly occurring structures (e.g., *Tf*+*Fn*, *Pg*+*Fa*) and previously uncharacterized assemblages.

To examine more intact, sessile biofilms, we also imaged SubP directly adherent to extracted tooth roots. This revealed additional previously uncharacterized microbial assemblages, including a distinct spherical structure termed “amorphous orbs”. Amorphous orbs are compact, near-spherical polymicrobial aggregates characterized by a rounded but irregular outer boundary and a stratified internal taxonomic organization, in which distinct taxa occupy spatially segregated concentric or layered regions (Fig. 1m–o; Extended Data Fig. 3s–z and aa–ag). Each orb comprised of a dense outer shell enriched in *Tf* and *Prevotella* sp., surrounding a central bacterial core. (Fig. 1n, o; Extended Data Fig. 3 t,u,w-z and ac-af). The morphology of the cells in the core suggested they may belong to poorly characterised Synergistetes cluster A taxa (e.g., *Fretibacterium*) for which reliable genus-level FISH probes are currently unavailable. Based on outer shell composition, orbs were classified into two subtypes: Type A (Fig. 1o; Extended Data Fig. 3ag), dominated by *Tf*, and *Fn*; and Type B (Fig. 1o; Extended Data Fig. 3a, g), lacking *Tf* but enriched in *Prevotella* and *Selenomonas*. Notably, the *Tf–Fn* co-localization observed in Type A orbs mirrors spatial interactions in TTBs.

### Polymicrobial TTBs exhibit distinct structural and taxonomic subtypes

We observed TTBs, a polymicrobial assemblage, in SubP from deep periodontal pockets (7 mm-13 mm). They represented some of the most conspicuous and morphologically distinct structures and were characterized by a long central filament (CF) core surrounded by orthogonally arranged shorter bristle-like bacterial cells (Fig. 2; Supplementary Movie 1). Single-channel hybridization with a universal bacterial probe (EUB338) highlighted the overarching architecture of TTBs (Fig. 2a, b; Supplementary Fig. S4 a-d), revealing CFs ranging from 20 to 120 µm in length and bristles arranged orthogonally along their axis. Species-specific FISH analysis identified *Fn* as a common constituent of the CF, while *Tf* frequently localized along the bristles (Fig. 2c; Supplementary Fig. S4 e,f; Supplementary Movie 2; Supplementary Fig. S5). In some instances, the bristles were co-colonized by *Prevotella sp*. (Fig. 2d; Supplementary Fig. S4 g,h, Supplementary Movie 3; Supplementary Fig. S6). Another type of TTBs observed were those with CFs not formed by *Fn* (Fig. 2f; Supplementary Fig. S4i). In one such configuration, the bristles were composed of *Prevotella sp.*, with members of Synergistetes cluster A surrounding the vicinity (Fig. 2f). In addition, the keystone pathogen *Pg* was observed in different planes of the Z-stacked TTBs, although it was rarely found closely integrated with the *Tf-Fn* core (Supplementary Fig. S4j.k).

**Fig. 2.**
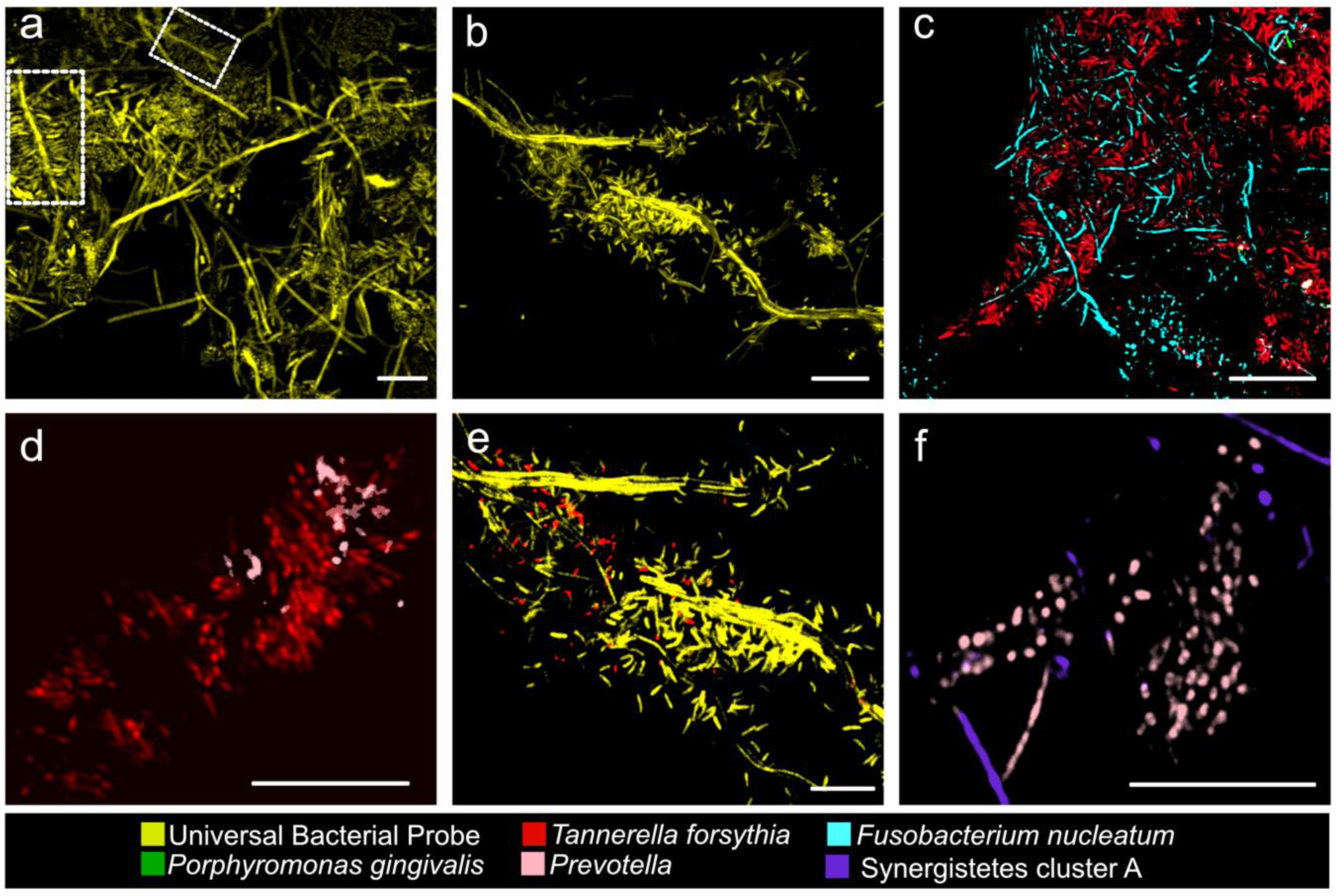
Spatial organization and species-level characterization of polymicrobial TTB structures in SubP from deep periodontal pockets (9 mm-13 mm). **a,b)** Single-channel hybridisation with the universal bacterial probe EUB338 highlights the TTB architecture. c) Fn commonly forms the CF, with *Tf* arranged as bristle-like projections. d,e) Bristles are occasionally co-colonized by *Prevotella*. f,g) Some TTBs exhibit CFs not formed by *Fn*, with *Prevotella*-dominated bristles and Synergistetes cluster A in the surrounding vicinity. Scale bars: 10 μm.

### TTB structural diversity, ecological composition, and clinical associations in subgingival plaque

To systematically categorize the diversity of TTBs observed in SubP, we developed a classification scheme based on total TTB length, CF morphology, and microbial composition (Supplementary Table 3). TTBs were broadly grouped into three size classes: macro (>60 µm), meso (40–60 µm), and micro (<40 µm), with each class further subdivided by CF architecture: either a single long filament or multiple short segments. This reflects the morphological continuum of TTBs, where all size classes could occur in either form, highlighting the structural and compositional plasticity of these polymicrobial assemblages.

We next evaluated whether TTB burden was associated with periodontal disease severity. Mixed-effects linear regression revealed that TTB density, measured as the number of TTB structures per field of view (FOV), was significantly associated with pocket depth, with each additional TTB per FOV corresponding to an estimated 2.43 mm increase in pocket depth (95% CI 0.78–4.08, p = 0.0038; Fig. 3a). Smoking status was not included in this model because only a single non-smoker was represented in the subset with available TTB/FOV measurements. To examine determinants of TTB occurrence across the broader cohort, mixed-effects logistic regression was used to model TTB presence at the site level. Pocket depth was a strong predictor of TTB detection, with each 1 mm increase in pocket depth associated with an approximately nine-fold increase in the odds of detecting TTBs (OR = 9.09, 95% CI 2.62–31.61, p = 0.00051; Fig. 3b, Supplementary Fig. S7). The fitted model indicated a sharp increase in TTB detection probability beyond approximately 9–10 mm pocket depth. (Fig. 3b). Despite local variation associated with pocket depth and surface type, normalized TTB density did not show obvious tooth-specific differences across sampled teeth (Fig. 3c). To examine broader spatial distribution patterns, we aggregated total detected TTB counts per tooth site. Teeth 26, 16, and 45 exhibited the highest observed counts (13, 12, and 10 TTBs, respectively). Although these values may also reflect differences in sampled plaque area and imaging coverage between sites.

**Fig. 3.**
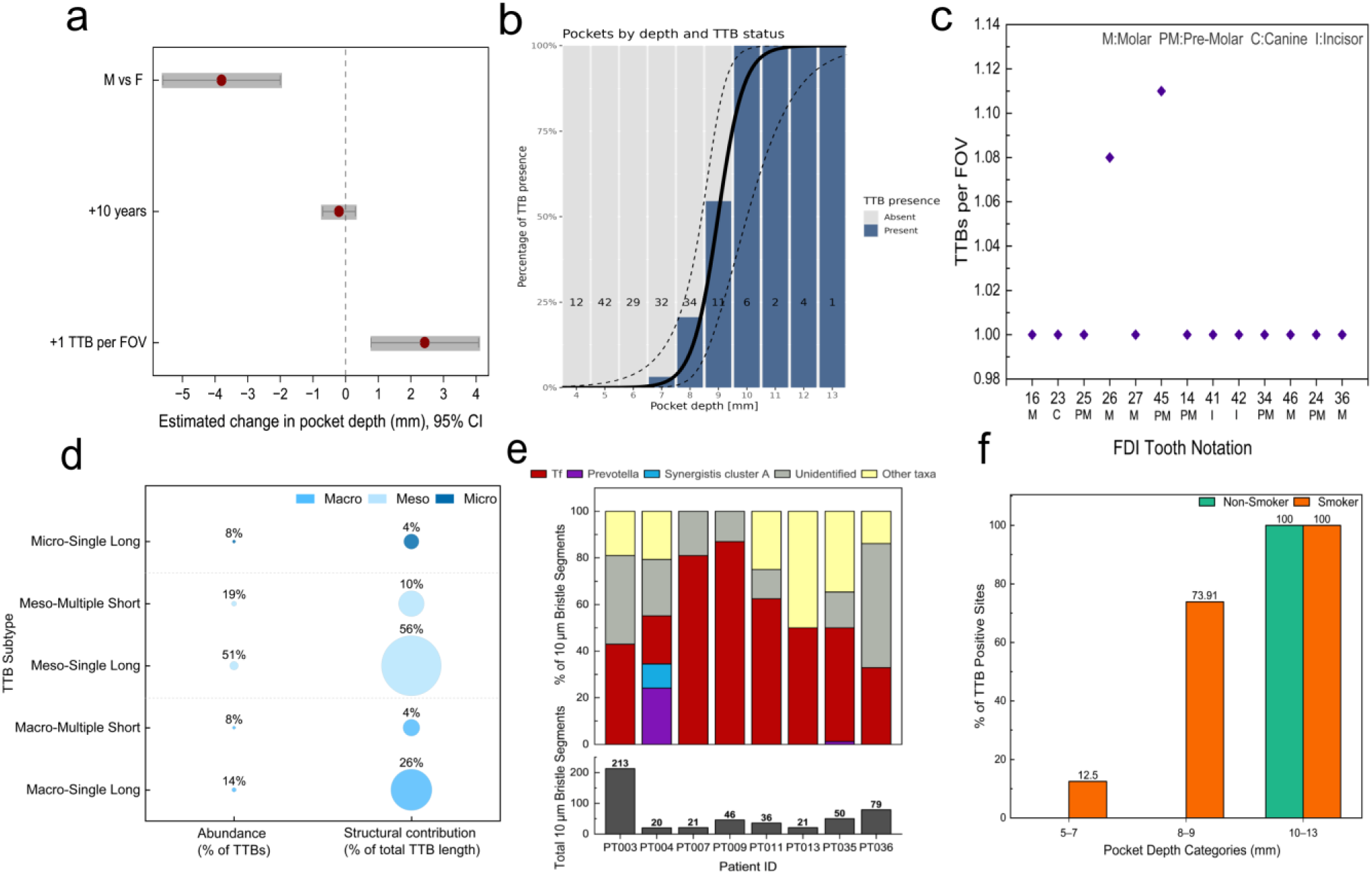
Structural and clinical correlates of Test Tube Brush (TTB) assemblages in subgingival plaque. a) TTB density is associated with increased pocket depth. This forest plot shows estimated changes in pocket depth from a mixed-effects linear regression model. Points represent estimates of model coefficients and horizontal lines indicate the corresponding 95% confidence intervals. The dashed vertical line indicates no estimated change. TTB density was positively associated with pocket depth, with each additional TTB structure per FOV corresponding to an estimated 2.43 mm increase in pocket depth. Age showed no clear association with pocket depth, while gender-associated differences were also observed (the mean pocket depths among female patients was 3.82mm larger than among the male patients from the cohort). b) Pocket depth predicts site-level TTB detection. Stacked bar plot showing the percentage of sampled subgingival sites with or without detectable test-tube brush (TTB) structures across recorded pocket depths. Bars represent site-level TTB status, with light grey indicating TTB-absent sites and blue indicating TTB-present sites. Numbers within bars indicate the total number of sampled sites at each pocket depth. The solid black curve shows the fitted mixed-effects logistic regression model predicting the probability of TTB detection as a function of pocket depth, and dashed curves indicate the 95% confidence interval. The model showed a sharp increase in TTB detection probability beyond approximately 9–10 mm pocket depth, consistent with pocket depth being a strong predictor of site-level TTB occurrence. c) Distribution of normalized TTB burden across the dentition. Scatter plot showing TTB abundance (TTBs per FOV) across individual teeth using FDI notation.). d) Bubble plot showing the relative abundance and estimated structural contribution of classified Test Tube Brush (TTB) subtypes. The left column shows each subtype as a percentage of total TTB abundance, while the right column shows its percentage contribution to total estimated cumulative TTB length. Bubble size is proportional to the corresponding percentage value. Total cumulative length was estimated by multiplying the number of TTBs in each subtype by the representative average length assigned to that subtype. Subtypes are coloured by structural size class: Macro, Meso, or Micro. e) Taxonomic composition of TTB bristle segments across patients. Top: Stacked bar plots showing the relative abundance of taxa within 10 µm bristle segments of TTB structures imaged from eight different patients (7 smokers and 1 non-smoker). Each bar represents the percentage contribution of each taxon (as identified by mFISH probes) to the total classified/segmented bristle regions per patient. Bottom: Total number of 10 µm bristle segments analyzed per patient, indicating imaging effort. f) Proportion of TTB-positive sites across pocket depth categories stratified by smoking status. Bar plot showing the percentage of periodontal sites positive for TTB structures across pocket depth categories (5–7 mm, 8–9 mm, and 10–13 mm) in smokers and non-smoker. Values above bars indicate the percentage of TTB-positive sites within each category. TTB prevalence increases with pocket depth in both groups, with smokers exhibiting TTB-positive sites at lower pocket depths compared to non-smoker.

Structural classification of 74 TTBs revealed that meso–single long TTBs were the dominant subtype, comprising 51.4% of structures and accounting for 55.6% of total cumulative TTB length (Fig. 3d, Supplementary Table 4). Macro–single long TTBs represented 13.5% of structures but contributed 26.3% of cumulative TTB length, reflecting their substantially greater representative filament length (∼90 µm). In contrast, macro–multiple short and micro–single long TTBs were comparatively rare, each representing 8.1% of structures, and contributed minimally to overall cumulative TTB length, accounting for 4.4% and 3.5%, respectively (Fig. 3d, Supplementary Table 4). Taxonomic profiling of TTB bristle segments highlighted compositional trends across patients. A total of 486 bristle segments (10 µm each) were analysed from eight patients. Because one smoker with a 5–7 mm periodontal pocket had no TTB-positive FOVs, the analysed cohort comprised seven smokers and one non-smoker. Across these segments, *T. forsythia* consistently dominated the outer bristles of TTBs (Fig. 3e). While other taxa such as *Prevotella*, Synergistetes cluster A, and unidentified organisms were variably present, *Tf* remained the most prevalent bristle-forming species regardless of patient or site.

Further, to analyse clinical associations, TTB presence was stratified by smoking status and pocket depth (Fig. 3f). TTB-positive sites were defined as sampled subgingival sites containing at least one detectable TTB structure. TTB-positive sites increased with pocket depth, rising from 12.5% in shallow pockets to 73.9% in intermediate pockets and reaching 100% in deeper pockets. Notably, TTBs were observed at intermediate pocket depths (8–9 mm) only in smoker-associated sites, whereas non-smoker-associated sites exhibited detectable TTBs only in deeper pockets (10–13 mm), suggesting earlier TTB emergence in smoker-associated disease (Fig. 3f).

### Structural remodeling of SubP: Transition from TTB to corncob assemblages post-treatment

To investigate architectural shifts in microbial communities and different assemblages following treatment, we revisited 18 subgingival sites from three selected patients (Pt_3, Pt_13, Pt_15) sampled both in 2023 and 2025. These patients, matched for age and gender, represented different smoking profiles and systemic conditions. In 2023, these sites harbored TTB structures within deep pockets with bleeding on probing (Extended Data Fig. 4).

Post-treatment imaging in 2025 revealed a complete loss of TTBs and the emergence of corncob-like assemblages largely localized to the biofilm periphery (Fig.4). Corncob-like structures have been reported earlier in healthy supragingival plaque and are considered hallmarks of oral health. Post-treatment corncobs consisted of a CF surrounded by radially arranged coccoid and rod-shaped bacteria. FISH identified that the inner core is dominated by *Lactobacillales* and *Bacillales*, with outer layers composed of Gammaproteobacteria and CFB cluster members. Interestingly, some core filaments failed to hybridize with universal or *Actinobacteria*-specific probes, suggesting the presence of probe-inaccessible or low-rRNA taxa as reported early.^12,13^

**Fig. 4.**
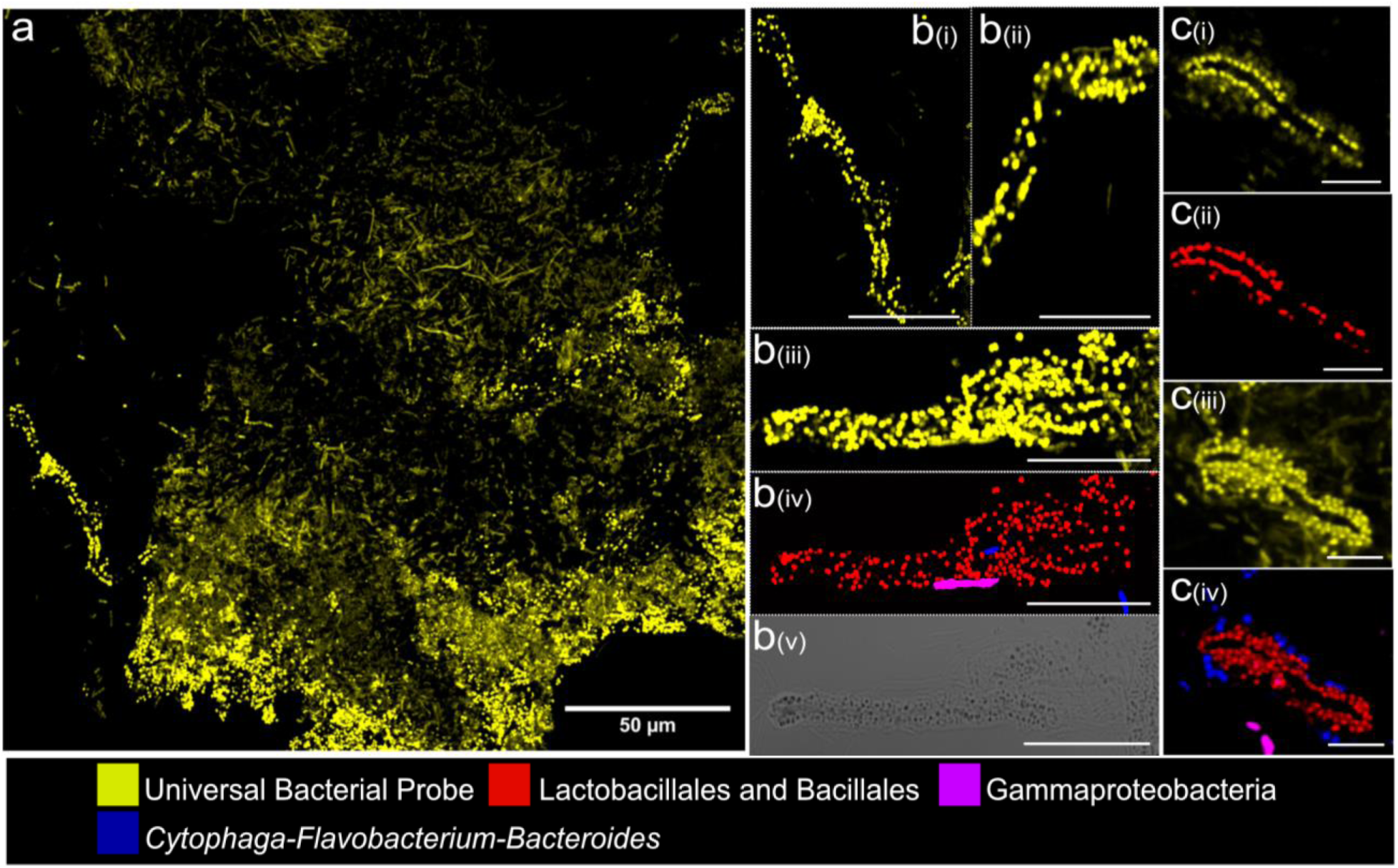
Longitudinal visualization of microbial biofilm architecture in selected patients’ post-treatment. **a)** Representative tile scan showing corncob assemblages located along the periphery of the subgingival biofilm, hybridized with a universal bacterial probe (EUB338). **b**) (i–iii) Single-channel image highlighting universal probe signal in representative corncobs. (iv) Taxonomic composition of corncob showing coccoid cells from the order Lactobacillales and Bacillales, filamentous members of Gammaproteobacteria, and sparsely distributed members of the CFB cluster (Cytophaga*–Flavobacterium–*Bacteroides,). (v) Bright-field image showing a central filament within the corncob that remained unstained by both the universal bacterial probe and the Actinobacteria-specific probe. **c)** Structural layering within corncobs: (i–ii) Initial layer comprising coccoid bacteria from Lactobacillales and Bacillales, overlaid by a second layer composed of CFB cluster bacteria and additional peripheral members include Gammaproteobacteria (iv). Scale bars: 50 µm a; 10 µm b–c.

Longitudinal imaging of 18 subgingival sites demonstrated a consistent post-treatment shift from TTB-dominated to corncob-dominated assemblages. To test whether this structural reorganization represented a directional treatment-associated transition, paired site-level architectural categories were compared before and after therapy using McNemar’s Chi-squared test with continuity correction. All nine TTB-positive sites transitioned to corncob-dominated architectures following treatment, indicating a significant directional shift among sites that were TTB-positive at baseline (McNemar’s χ² = 7.11, p = 0.0077). The consistent emergence of corncob structures across all three patients suggests a convergent post-treatment remodeling pattern irrespective of smoking or diabetic status.

## Discussion

Microbiome research has transformed our understanding of human disease by defining microbial taxa and functional pathways associated with health and pathology. However, for biofilm-associated microbiomes, sequencing-based approaches necessarily disrupt community structure and therefore cannot resolve the spatial organisation through which microorganisms interact *in vivo*^16^. By preserving and resolving the taxon-resolved architecture of the human subgingival plaque microbiome across clinically characterised periodontal sites, we show that disease-associated biofilms are organised into recurrent multicellular assemblages whose taxonomic composition, geometry and clinical distribution are reproducible across individuals. These findings identify microbial architecture as a biologically meaningful organisational layer of the microbiome that complements community composition and functional potential, providing a previously inaccessible axis for interpreting microbiome-associated disease. Collectively, our findings support four conceptual advances that redefine how disease-associated biofilm microbiomes can be interpreted (Figure 5). Microbiome research has transformed our understanding of human disease by defining microbial taxa and functional pathways associated with health and pathology. However, for biofilm-associated microbiomes, sequencing-based approaches necessarily disrupt community structure and therefore cannot resolve the spatial organisation through which microorganisms interact *in vivo*^16^. By preserving and resolving the taxon-resolved architecture of the human subgingival plaque microbiome across clinically characterised periodontal sites, we show that disease-associated biofilms are organised into recurrent multicellular assemblages whose taxonomic composition, geometry and clinical distribution are reproducible across individuals. These findings identify microbial architecture as a biologically meaningful organisational layer of the microbiome that complements community composition and functional potential, providing a previously inaccessible axis for interpreting microbiome-associated disease. Collectively, our findings support four conceptual advances that redefine how disease-associated biofilm microbiomes can be interpreted.

**Fig. 5.**
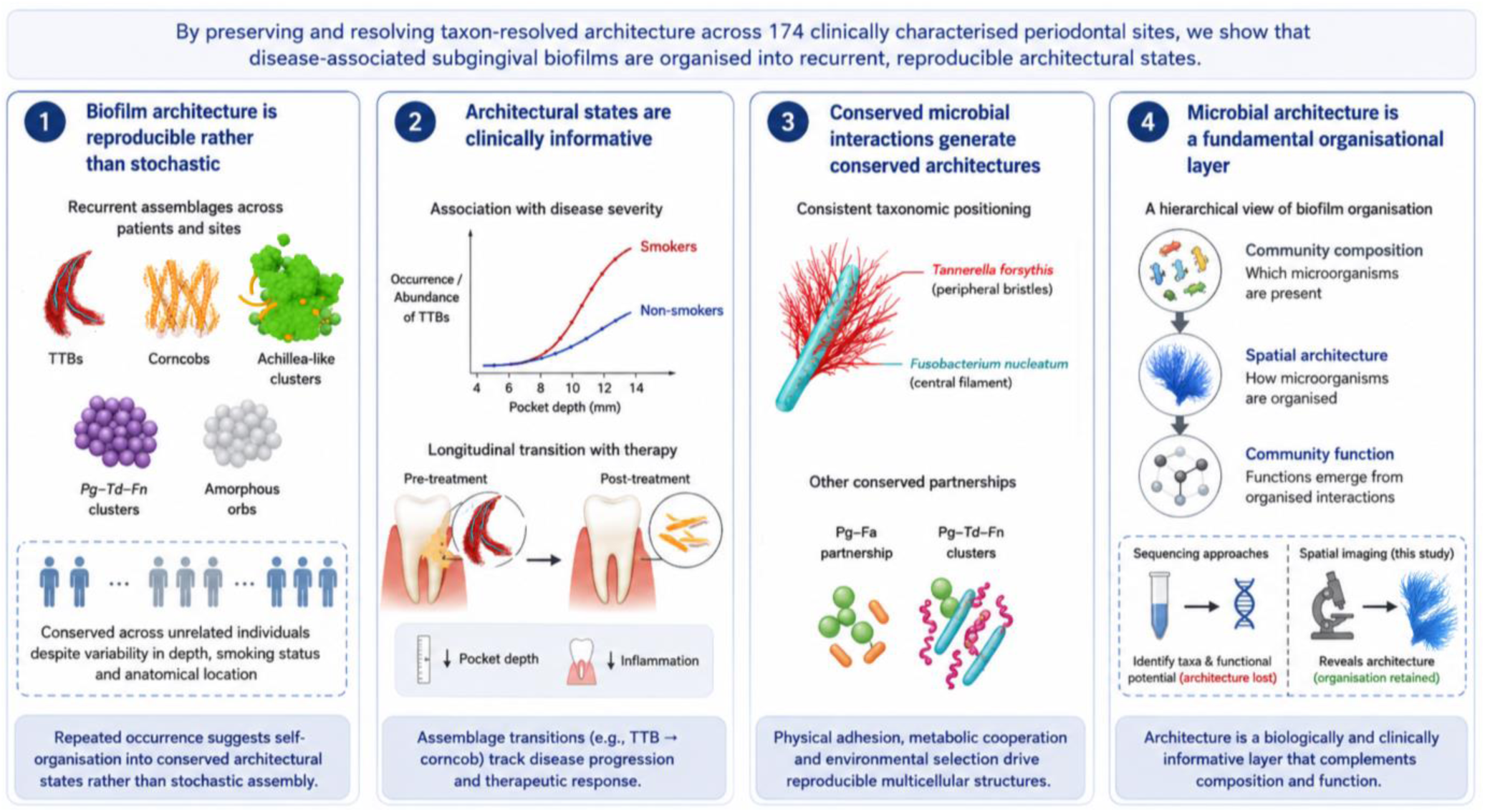
Conceptual model of disease-associated biofilms as structured and responsive ecosystems. Spatial organisation of SubP is reproducible rather than stochastic, with conserved microbial interactions generating recurrent architectural states. These states are clinically informative, varying with disease severity, smoking and periodontal therapy. Together, the findings position microbial architecture as a fundamental organisational layer of biofilm-associated microbiomes that complements community composition and function.

### Biofilm architecture is reproducible rather than stochastic

A central finding of this study is that spatial architecture follows reproducible organisational rules. Across 174 SubP samples from clinically characterised periodontal sites, canonical oral architectures, including test-tube brushes (TTBs) and corncobs, coexisted with previously undescribed assemblages such as *Achillea*-like clusters, *Pg–Td–Fn* clusters and amorphous orbs. Despite substantial heterogeneity in pocket depth, smoking status and anatomical location, these assemblages repeatedly exhibited conserved taxonomic organisation across unrelated individuals. Such recurrence argues against stochastic spatial assembly and instead suggests that microbial communities self-organise into conserved architectural states. We propose that these architectures represent stable ecological solutions to the physicochemical constraints of the periodontal pocket, analogous to the way environmental selection repeatedly shapes community composition across larger ecological scales.

### Architectural states are clinically informative

The observed architectures were not distributed uniformly throughout the cohort but occupied distinct ecological niches. Among all assemblages, TTBs showed the strongest association with disease severity, with both their occurrence and abundance increasing with pocket depth and their emergence occurring at shallower depths in smokers. In contrast, other architectures occurred across broader clinical contexts, suggesting differences in ecological breadth. Longitudinal sampling further demonstrated a reproducible transition from TTB-dominated to corncob-dominated biofilms following periodontal therapy, concomitant with reductions in pocket depth and resolution of disease. Hence, these observations indicate that disease progression and therapeutic response involve transitions between reproducible architectural states rather than simply changes in microbial composition or biomass. Within this framework, TTBs are interpreted not merely as distinctive morphologies but as spatial signatures of advanced ecological states within the periodontal pocket.

### Conserved microbial interactions generate conserved architectures

The repeated organisation of specific taxa within recurrent architectures suggests that spatial assembly is governed by conserved microbial interactions. Across patients, *Fusobacterium nucleatum* consistently formed the central filament of TTBs, whereas *Tannerella forsythia* repeatedly occupied the peripheral bristles. Similar taxonomic conservation characterised other recurrent assemblages, including *Pg–Fa* and *Pg–Td–Fn* partnerships. Previous studies have demonstrated coaggregation and metabolic cooperation between several of these organisms *in vitro^17–19^*, here, we demonstrate that these interactions repeatedly organise into defined three-dimensional architectures *in vivo*. This conservation implies that microbial spatial organisation is constrained by ecological interactions rather than species co-occurrence alone, integrating physical adhesion, metabolic cooperation and local environmental selection into reproducible multicellular structures.

### Microbial architecture represents a fundamental organisational layer of biofilm-associated microbiomes

Collectively, these findings support a hierarchical view of microbiome organisation in which community composition defines which microorganisms are present, spatial architecture defines how those microorganisms are organised, and community function emerges from these organised interactions. These organisational layers are complementary rather than interchangeable. Sequencing has transformed our ability to identify community members and predict functional potential but cannot recover the architectural layer because spatial relationships are lost during sample processing. This framework identifies microbial architecture as a fundamental organisational layer of biofilm-associated microbiomes. Within this framework, disease-associated biofilms are interpreted not simply as dysbiotic communities but as ecosystems occupying discrete architectural states whose organisation is itself biologically and clinically informative.

While this study establishes an architectural framework for interpreting disease-associated biofilms, several important questions remain. Because our analyses are observational, they identify conserved architectural states but cannot determine the mechanisms by which these structures arise or whether they actively contribute to disease progression. Likewise, although multiplex FISH enabled taxon-resolved imaging of key periodontal organisms, some microorganisms remained taxonomically unresolved, highlighting the need for expanded probe libraries and complementary spatial omics approaches. Finally, although periodontitis provides a tractable model for investigating host-associated biofilms, whether similar architectural states occur across other biofilm-associated microbiomes remains unknown. Addressing these questions will require integrating high-resolution spatial imaging with experimentally tractable biofilm models, spatial transcriptomics, metabolomics, perturbation-based experiments and quantitative ecological modelling to determine how architectural states develop, remodel during disease progression and treatment, and influence microbial community function.

Although this work focuses on periodontitis, the implications extend beyond the oral cavity. Host-associated biofilms throughout the body are similarly shaped by local physicochemical gradients, attachment surfaces and interspecies interactions, suggesting that reproducible architectural states may represent a general organisational principle of biofilm-associated microbiomes. Our findings provide experimental evidence that spatial architecture is not simply a descriptive property of microbial communities but a fundamental organisational dimension that complements community composition and function. More broadly, they establish a framework for investigating how microbial architecture contributes to biofilm-associated disease and provide a foundation for developing future architecture-informed diagnostics and therapeutic monitoring strategies.

## Methods

### Patient Recruitment and Clinical Assessment

Patients with a clinical diagnosis of periodontitis were recruited under institutional ethics approval (2023-26641-41275-3). Inclusion criteria required the presence of at least one periodontal site with a probing depth ≥ 5 mm. Exclusion criteria included recent antibiotic use, pregnancy, or systemic inflammatory diseases (except diabetes). All participants provided informed consent. Clinical assessments were conducted by an experienced periodontist and included probing depth, bleeding or suppuration on probing, gingival inflammation, smoking status (non-smoker, light smoker [<10 cigarettes/day], or heavy smoker [≥10 cigarettes/day]), age, gender, and comorbidities. These data were recorded and used for downstream multivariate analyses and cohort stratification (Supplementary Table 1). The tooth-site nomenclature used in Supplementary Table 1, Extended Data Fig. 1a, and throughout the manuscript follows the FDI World Dental Federation notation (ISO 3950: *Dentistry—Designation system for teeth and areas of the oral cavity*)^20^.

SubP samples were collected using a single stroke of a sterile curette from sites exhibiting deep periodontal pockets (typically >5 mm) in patients classified as Stage III or IV according to the 2017 World Workshop classification ^5,21^. Samples were placed into sterile Oral Bacterial Growth Medium (OBGM) and stored at 4 °C for up to 1 hour before fixation. Extracted teeth were also obtained in situations where the removal of a very mobile tooth was required as part of the patient’s treatment. For SubP sampled from extracted teeth, the gingival margin was marked prior to extraction, and teeth were removed with minimum trauma by gripping only the crown to avoid disturbing the subgingival biofilm. No instrumentation was introduced subgingivally. The location of plaque deposits was recorded relative to the gingival margin and the deepest point of observed biofilm, corresponding to the pocket base.

### Subgingival Plaque Sampling and Fixation

SubP samples were collected from three regions: (1) deep periodontal pockets and (2) shallow pockets post treatment (3) root of extracted tooth. Sampling from periodontal pockets was performed using sterile curettes under isolation, targeting the deepest sites with visible inflammation. Tooth-attached SubP was recovered by gentle scraping of exposed root surfaces. Samples were immediately fixed in 2% paraformaldehyde in PBS for 4 hours on ice and stored in 1:1 10mM Tric HCl (pH 7.5): 96% ethanol at −20°C until processing.

### Fluorescence *In Situ* Hybridization (FISH)

FISH was performed by modifying standard protocols. ^22^ PFA-fixed SubP samples (100μl) in 1:1 20mM Tris HCl (pH 7.5): 50% ethanol were transferred onto <u>Poly-L-lysine–coated glass slides</u> and dried at 46 °C for 30 minutes. Slides were pre-incubated in hybridization buffer (0.9 M NaCl, 20 mM Tris-HCl, pH 7.5) for 15 min at 46 °C. Hybridization was performed using 2 mM of each probe in hybridization buffer (900 mM NaCl, 20 mM Tris-HCl, pH 7.5, 0.02% SDS, 30% formamide) in a humid chamber for 4-5h at 46 °C. Slides were washed thrice in wash buffer (215 mM NaCl, 20 mM Tris-HCl, 5 mM EDTA, pH 7.5) at 48 °C for 10 min each, rinsed in ice-cold water, and air-dried. Mounting was done in a 60% glycerol solution. For pure cultures, 10–20 μl of fixed culture was hybridized with 1 mM of probe under identical conditions.

Fluorophore-labeled oligonucleotide probes were purchased from <u>ThermoFisher</u> or <u>biomers.net</u> and validated against pure cultures of respective bacteria (Supplementary Fig. S8). A full list of taxonomic targets, probe sequences, fluorophore tags and probe sets are provided in Supplementary Table 5 and 6 respectively.

### Spectral/Confocal and Airyscan Imaging

Imaging was performed on a ZEISS LSM 980 with Airyscan 2 and a ZEISS LSM 880 Airyscan Fast confocal microscopes. The LSM 980 was equipped with 405, 445, 488, 515, 561, 594, 639, and 730 nm laser lines with two photomultiplier tube (PMT), 32 channel gallium arsenide phosphide (GaAsP) spectral array detector, near infrared (NIR) GaAs detector and a 32 channel GaAsP array Airyscan detector. The LSM 880 was equipped with 405, 458, 488, 514, 561, 594, and 633 nm laser lines, with the same detector configuration as the LSM980 Airyscan 2, except without the NIR-GaAs detector. For imaging, whole plaque tile scans were acquired using the 25×/0.8 NA Plan-Apochromat lens (Item number 420852-9871-000), and the high magnification images were acquired using the 40×/1.3 NA Plan-Apochromat lens (Item number 421762-9900-799), or the 63×/1.4 NA Plan-Apochromat lens (Item number 421782-9900-799). Most datasets acquired in standard confocal mode were acquired with pixel sizes set to yield 1.2x Nyquist sampling rate and scanned sequentially per channel to minimize bleed-through. Spectral imaging was performed on selected samples using lambda scanning mode to capture the full spectral emission profiles, followed by linear unmixing in ZEN Blue 3.6 or Zen Black 2.3 software using reference spectra from the single-probe controls. Probe concentrations were empirically adjusted to improve the signal-to-noise ratio in the image. For Airyscan imaging, the pixel sizes were set at 2x the Nyquist criterion and images were processed using standard Airyscan processing module on Zen Blue 3.6 or Zen Black 2.3 software. Image processing and analysis, including maximum intensity projections, channel alignment, and figure assembly, were conducted in Fiji (ImageJ) ^23^.

### Custom Fiji Macro Development and Image Analysis Workflow

To ensure reproducible and consistent image handling across large datasets, we developed custom Fiji (ImageJ)^23^ macros for semi-automated batch processing and quantitative analysis. These scripts were authored specifically for this study to streamline repetitive processing steps and to enforce uniform application of adjustments across all images within a dataset.

The batch-processing macros were applied wherever possible to automate steps such as channel splitting/merging, maximum intensity projection generation, channel alignment, linear brightness/contrast adjustment, and scale bar placement. These macros were designed to process entire folders of raw .czi files exported from Zen Blue 3.6 or Zen Black 2.3, applying identical parameters to each image to maintain consistency and minimise user bias.

For datasets requiring figure-specific processing (e.g., unusual probe combinations, rare structures, or atypical image sizes), tailored macro scripts were written to accommodate these unique conditions. These figure-specific macros retained the same principles of reproducibility: adjustments were always linear and applied equally across channels, and no nonlinear contrast enhancements or selective alterations were performed. All custom Fiji macro scripts developed in this study for semi-automated batch processing of TTBs, figure-specific image adjustments, and quantitative analysis are currently under private embargo will be submitted and made public upon request. A DOI and direct access link will be provided in the final version of the article.

### TTB Classification and Taxonomic Composition Analysis

TTBs were defined as filamentous microbial assemblages with a clearly identifiable central filament and radially arranged bristle-like peripheral cells, extending ≥20 μm in length. For each of eight patients, 10 fields of view (FOVs) were acquired per probe set, selected based on high signal-to-noise ratio, structural integrity of TTBs, and consistent multi-channel hybridization.

Each TTB was divided longitudinally into contiguous 10 μm units for compositional analysis. Segment units were included only when both core and peripheral architecture were well resolved across the z-stack. Within each segment unit, bacterial identity was determined using genus- or species-specific FISH probes. Segments were categorized into single-taxon (e.g., *Tf*), dual-taxon (e.g., *Tf+Prevotella*), or multi-taxon configurations (e.g., *Tf+Prevotella +* Synergistetes cluster A). Segments that lacked probe signal but retained TTB morphology were classified as uncharacterized. A total of 486 segments were quantified across patients. Segment identities were compiled into a taxonomic matrix and visualized as stacked bar plots to show compositional variability (Fig. 3e top panel), and patient-level bar charts were generated to display the number of segments analyzed per case (Fig. 3e bottom panel). Extended data figure 5 represents the detailed workflow for TTB quantification with an example.

### Statistical Analysis, Data Visualisation and Plotting

Clinical and compositional data were analyzed and visualized using <u>Python 3.11</u> (<u>Pandas,</u> <u>Plotly</u> <u>v5.0+</u>). These data were exported as SVGs and post-processed in <u>Affinity Photo 2.5.2</u> and <u>Affinity</u> <u>Designer 2.6.5</u> for color annotation and layout standardization. Additional were plotted in <u>Origin Pro</u> <u>20205 (10.2)</u>.

Mixed-effects regression models were fitted to account for repeated measurements and multiple sampling sites within individual patients. Four mixed-effects models were fitted. First, a logistic mixed-effects model was used to assess whether bleeding on probing was associated with pocket depth, age, gender, tooth group and smoking status. Second, a linear mixed-effects model was used to assess associations between pocket depth and age, gender, tooth group and smoking status. Third, a linear mixed-effects model was fitted to evaluate the association between pocket depth and TTB density, expressed as the number of TTBs per field of view, while adjusting for age and gender. Finally, a logistic mixed-effects model was used to assess whether TTB presence was associated with pocket depth, age, tooth group and smoking status.

For continuous and dichotomous predictors, Wald Z-tests were used to assess whether model coefficients differed significantly from zero. For the categorical tooth-group predictor, Type II Wald chi-squared tests were used to assess the overall association between tooth group and the response variable. Where a significant overall association was detected, post-hoc comparisons between estimated marginal means were performed, with p-values adjusted for multiple comparisons using Tukey’s method.

Mixed-effects analyses were conducted in the R Statistical Computing Environment (v4.5.1) using the "glmmTMB" package ^24^. Estimated marginal means and post-hoc comparisons were calculated using the "emmeans" package ^25^. Other statistical analyses were performed using either in <u>Origin Pro 20205</u> <u>(10.2)</u> or <u>SciPy v1.11.4 Python 3.11</u>. A summary of all statistical analyses, including data level, sample size, outcome variables, predictors and statistical tests or models, is provided in Supplementary Table 8.

## Acknowledgements

This project was supported by the Australian National Health and Medical Research Council grants to D.G. (Grant ID: APP1196924) and E.C.R. (Grant IDs: APP1193647 and APP1123866). DG is supported by a Cumming Global Centre for Pandemic Therapeutics Foundation Grant (CGCPT00060). AP is supported by the Melbourne Research Scholarship and Rowden White Scholarship. We acknowledge the Biological Optical Microscopy Platform (University of Melbourne) and the staff Dr. Gabriela Segal Wasserman and Dr. Elie Cho for their advice on image processing for access to microscopy facilities. During manuscript preparation, large language model tools were used to assist with structural refinement of text. All scientific content, analyses, interpretations and conclusions were generated, verified and approved by the authors.

## Authors contributions

Conceptualization: P.D.V., E.C.R., D.G.; study design, P.D.V., A.P., E.C.R., D.G.; methodology: A.P., P.M., H.C., H.P., P.D.V.; Sample collection: L.N., A.P., S.B.S; formal analysis: P.D.V.; A.P.; D.G; E.C.R; S.G.D.; investigation, A.P.; P.D.V., E.C.R. D.G.; image analysis and FIJI codes: H.P., A.P; statistical planning and analysis, A.P., J.S. E.C.R., D.G.; data curation, A.P.; writing original draft, AP.; review & editing, all authors; funding acquisition, E.C.R., D.G.

## Data availability

The data supporting the findings of this study are included within the article and its Supplementary Information files. Additional source data and image-analysis outputs will be made available through an appropriate public or institutional repository upon publication.

## Code availability

Custom Fiji/ImageJ macros and scripts, Python code, and R code used for image processing, data analysis, statistical modelling and visualisation will be deposited in an appropriate public or institutional repository upon publication.

