## Supplementary material for "The Spatial Architecture of Human Subgingival Biofilms": ExtendedData_SubP.pdf

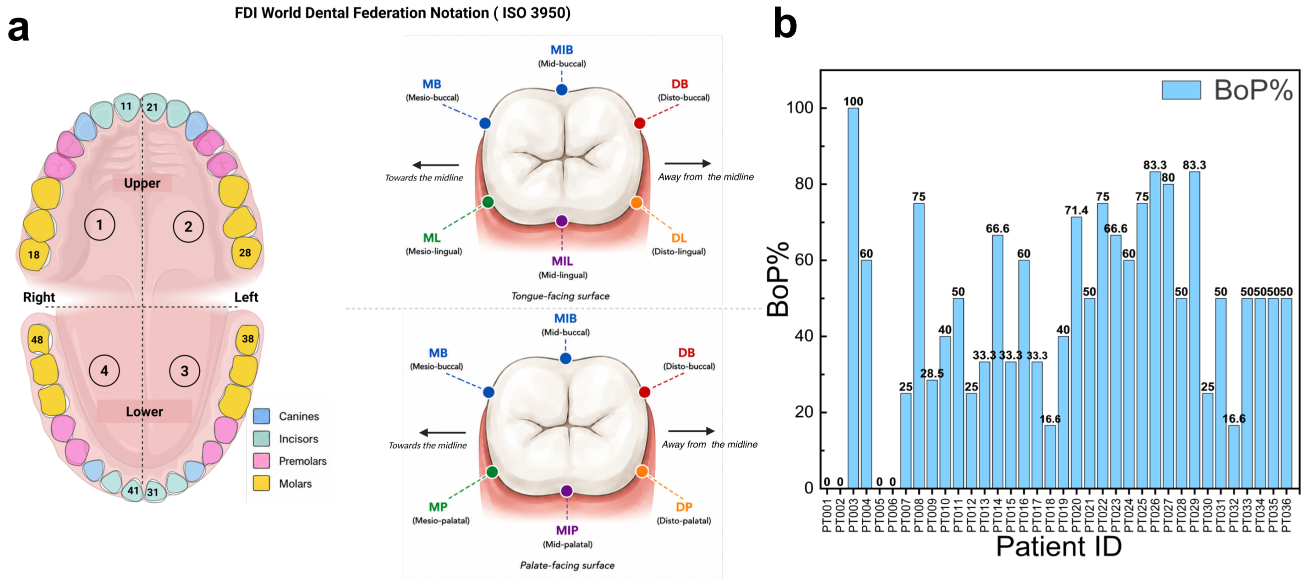


**Extended Data Fig. 1| a,** Schematic showing the FDI World Dental Federation tooth-numbering system and anatomical surface orientation used for site-level annotation of SubP samples. Quadrants are indicated using standard FDI notation, with representative tooth numbers shown for central incisors 11, 21, 41 and 31 and third molars 18, 28, 48 and 38. Tooth classes are colour-coded as incisors, canines, premolars and molars. Each sampled site is designated by a tooth number followed by a surface abbreviation: MB, mesiobuccal; MIB, midbuccal; DB, distobuccal; ML, mesiolingual; MIL, midlingual (mandibular teeth); and DL, distolingual. For maxillary teeth, lingual sites are replaced by their palatal equivalents (MP, mesiopalatal; MIP, midpalatal; and DP, distopalatal). This nomenclature is used consistently throughout the manuscript and Supplementary Table 1 to identify all subgingival sampling sites. **b.** Percentage of BoP-positive sites per patient. BoP% was calculated as (BoP-positive sites/total sites) ×100 for each patient. Patients are shown in the same order as in the Max PD plot. Spearman's correlation showed a weak, non-significant association between Max PD and BoP % ($\rho$ = 0.24, p = 0.16), indicating BoP is not a reliable marker of disease severity in this cohort.

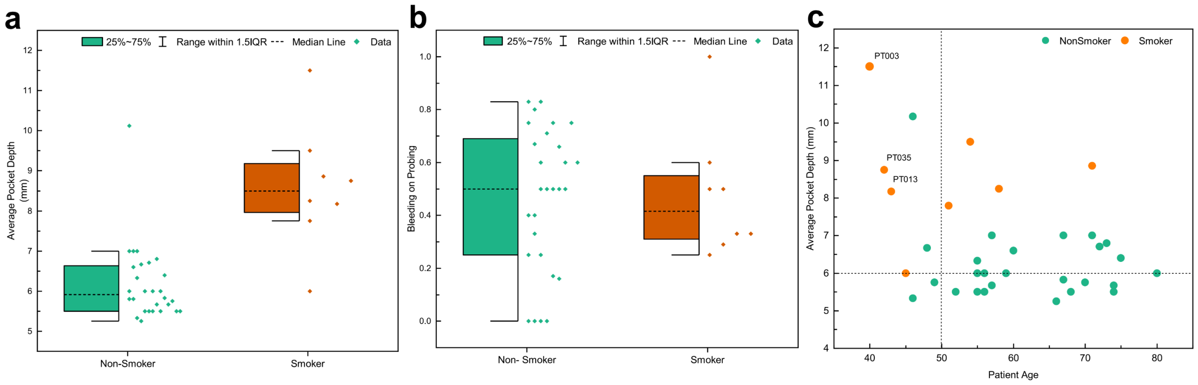


**Extended Data Fig. 2. | Smoking status and age influence periodontal pocket depth and site-specific disease patterns. a, b**) Smokers exhibit significantly deeper periodontal pockets but comparable levels of bleeding on probing (BoP) relative to non-smokers. **a**) Boxplots comparing average pocket depth per patient between smokers (n = 8) and non-smokers (n = 28). Boxes represent the interquartile range (25%–75%), whiskers span 1.5× IQR, and medians are marked by dashed lines; individual data points presented to the right of the boxes. Smokers exhibited significantly greater pocket depths than non-smokers ($p < 0.0001$, Mann–Whitney U test). **b**) Boxplots display the proportion of BoP-positive sites per patient in smokers and non-smokers. No significant difference in BoP levels was observed between groups ($p = 0.86$, Mann–Whitney U test). **c**) Smokers under 50 years of age exhibit elevated periodontal pocket depths. Scatter plot showing mean pocket depth per patient as a function of age, with individuals colored by smoking status (teal= non-smoker, orange = smoker). Dashed lines indicate the clinical pocket-depth threshold of 6 mm (horizontal) and an age reference cutoff of 50 years (vertical). Several smokers under 50 years of age (PT003, PT013, and PT035) exhibited mean pocket depths exceeding 8 mm, whereas most non-smokers across age groups clustered near or below the 6 mm threshold except for PT036. These trends suggest an association between smoking and increased periodontal disease severity independent of age distribution within the cohort.

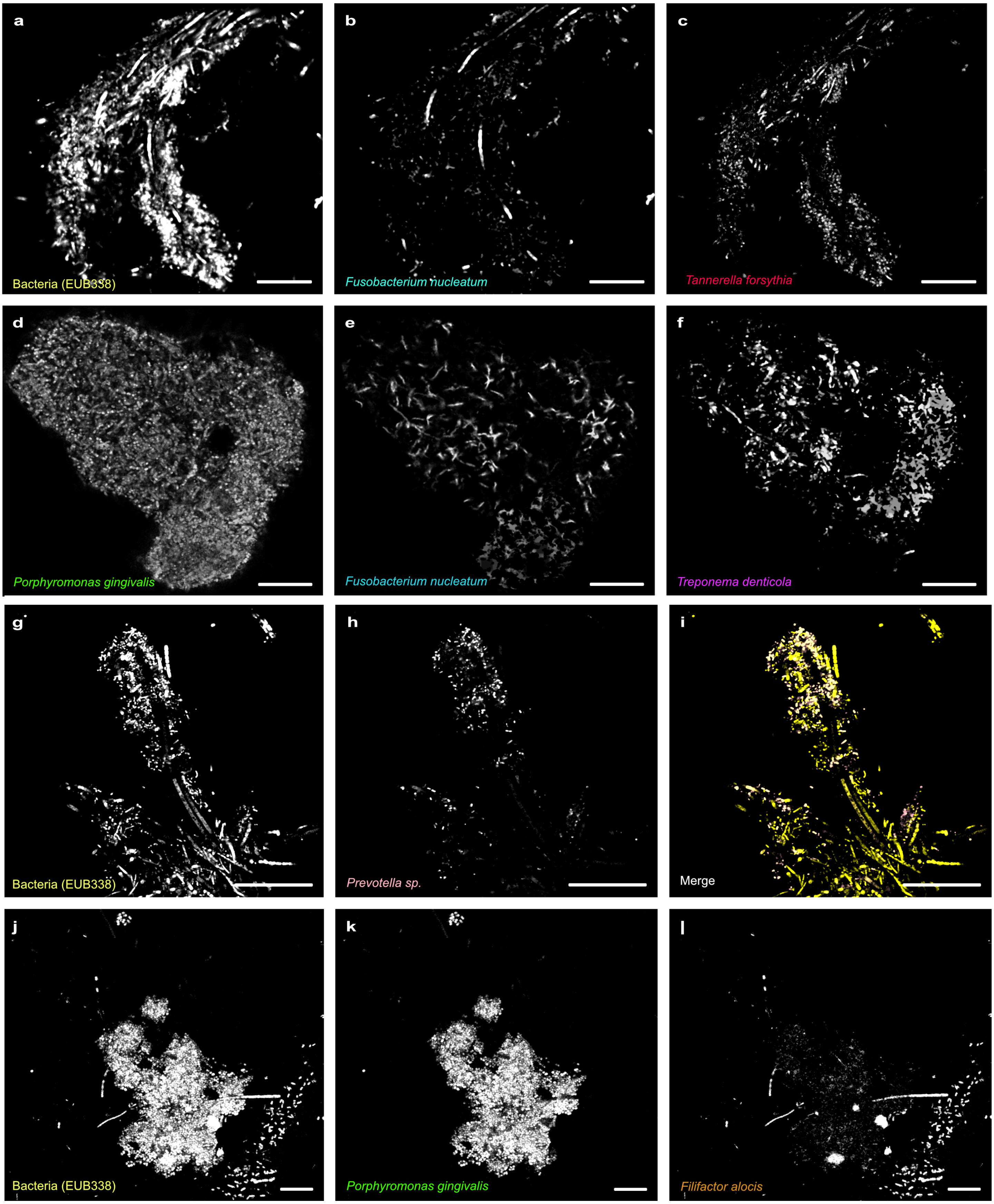
a
Bacteria (EUB338)
b
Fusobacterium nucleatum
c
Tannerella forsythia
d
Porphyromonas gingivalis
e
Fusobacterium nucleatum
f
Treponema denticola
g
Bacteria (EUB338)
h
Prevotella sp.
i
Merge
j
Bacteria (EUB338)
k
Porphyromonas gingivalis
l
Filifactor alocis

47

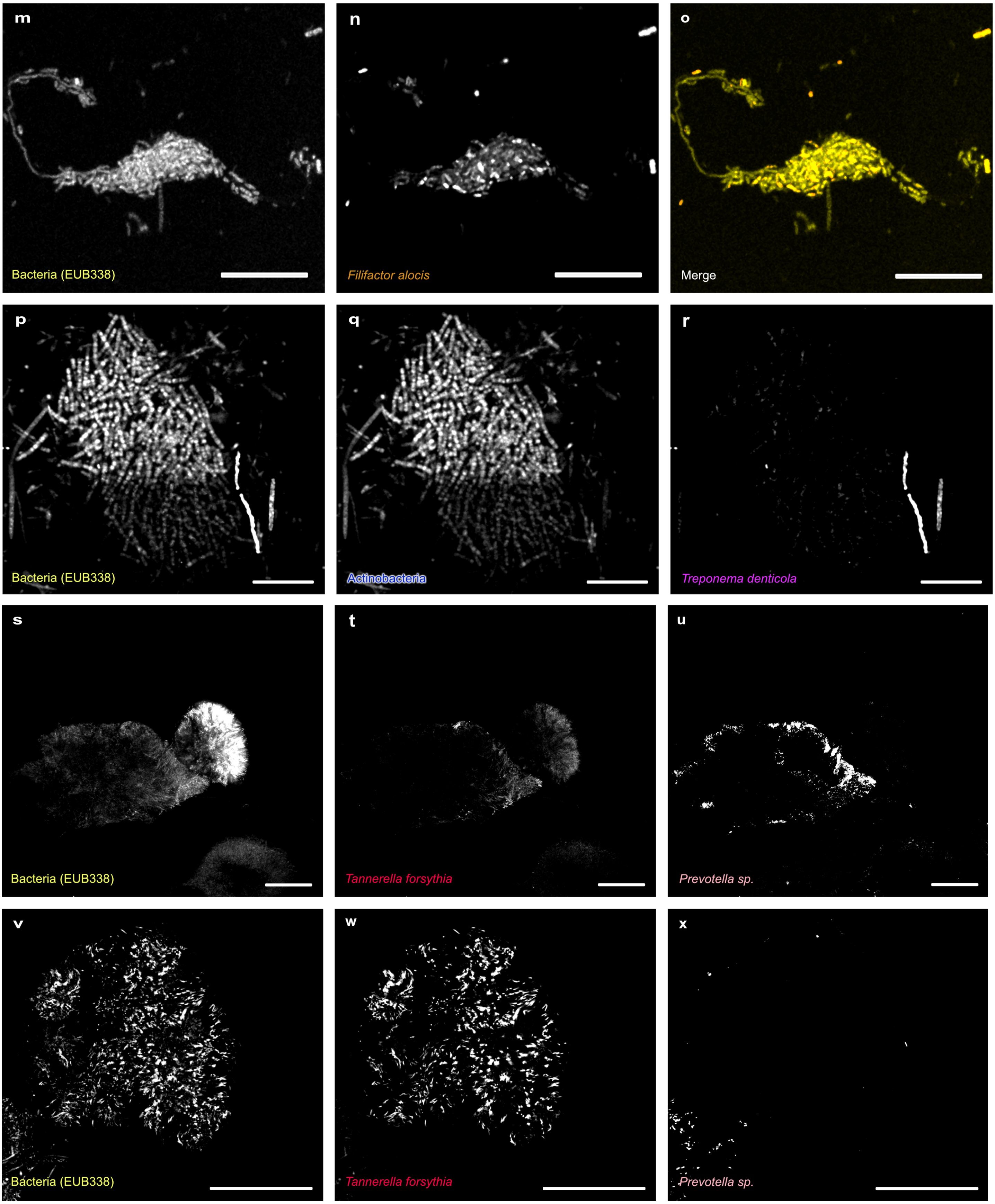
m
Bacteria (EUB338)
n
Filifactor alocis
o
Merge
p
Bacteria (EUB338)
q
Actinobacteria
r
Treponema denticola
s
Bacteria (EUB338)
t
Tannerella forsythia
u
Prevotella sp.
v
Bacteria (EUB338)
w
Tannerella forsythia
x
Prevotella sp.

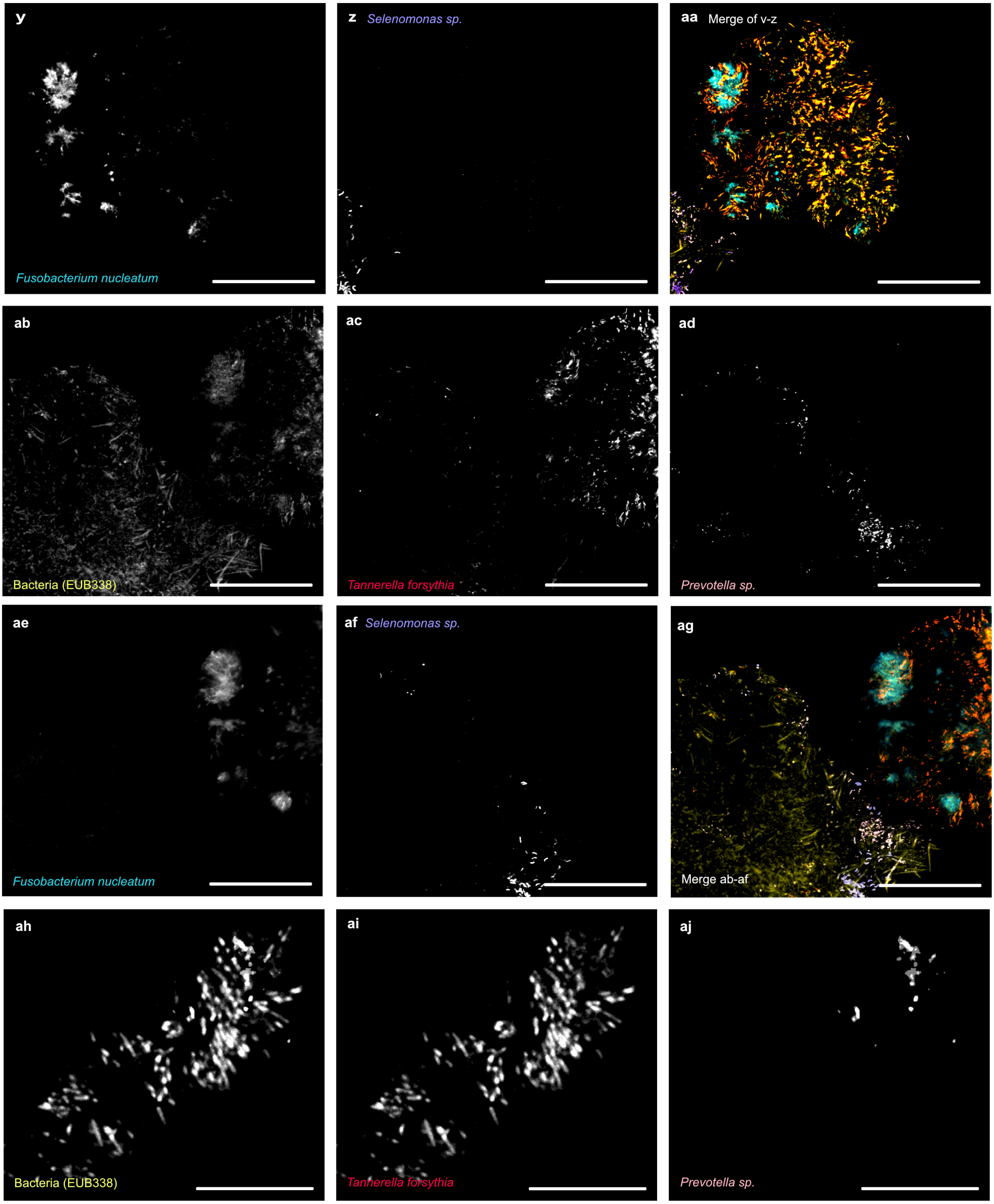

y
Fusobacterium nucleatum
z
Selenomonas sp.
aa
Merge of v-z
ab
Bacteria (EUB338)
ac
Tannerella forsythia
ad
Prevotella sp.
ae
Fusobacterium nucleatum
af
Selenomonas sp.
ag
Merge ab-af
ah
Bacteria (EUB338)
ai
Tannerella forsythia
aj
Prevotella sp.


50

**Extended Data Fig. 3. | Grayscale reference images for multiplexed FISH channels.** Representative grayscale images corresponding to the individual fluorescence channels shown in various main figures. These panels illustrate structures in which multiple bacterial probes either co-localize or are superimposed within the same spatial region, making them indistinguishable in merged images shown in the main figures in the article. Each grayscale image highlights a single probe channel, with text labels in the corresponding pseudocolour used in the article to aid interpretation (for example, a = EUB338, labelled in yellow; b = *Fn*, labelled in cyan; c = *Tf*, labelled in red). Certain panels for like (g–i) depict two-species assemblages, where (g) shows EUB338 (yellow), (h) shows *Prevotella sp.*(pink), and (i) presents their merged image in colour. Scale bars:10 µm.

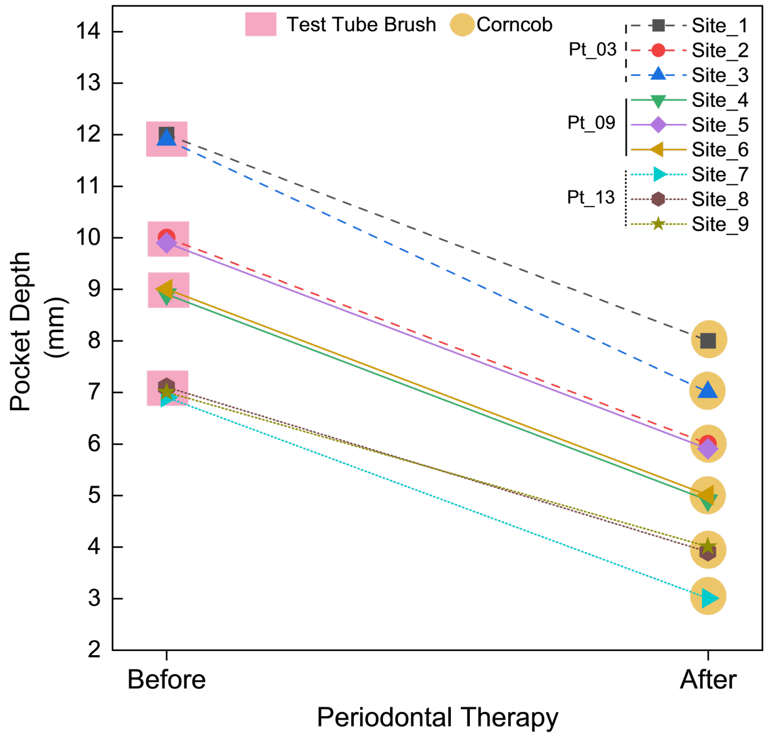


**Extended Data Fig. 4. | Longitudinal reduction in pocket depth and associated structural transition across nine subgingival sites following periodontal treatment.** Paired line plot showing changes in pocket depth before and after treatment across nine matched subgingival sites sampled from three patients (Pt_3, Pt_09 and Pt_13; n = 9 sites total, 3 sites per patient). Each line represents an individual site tracked longitudinally. All sites showed a reduction in pocket depth following therapy, accompanied by a complete structural transition from test tube brushes (pink squares) pre-treatment to corncobs (golden squares) post-treatment. Structural transition between pre- and post-treatment states was significant (McNemar's $\chi^2 = 7.11$, $p = 0.0077$), supporting non-random architectural remodeling following treatment.

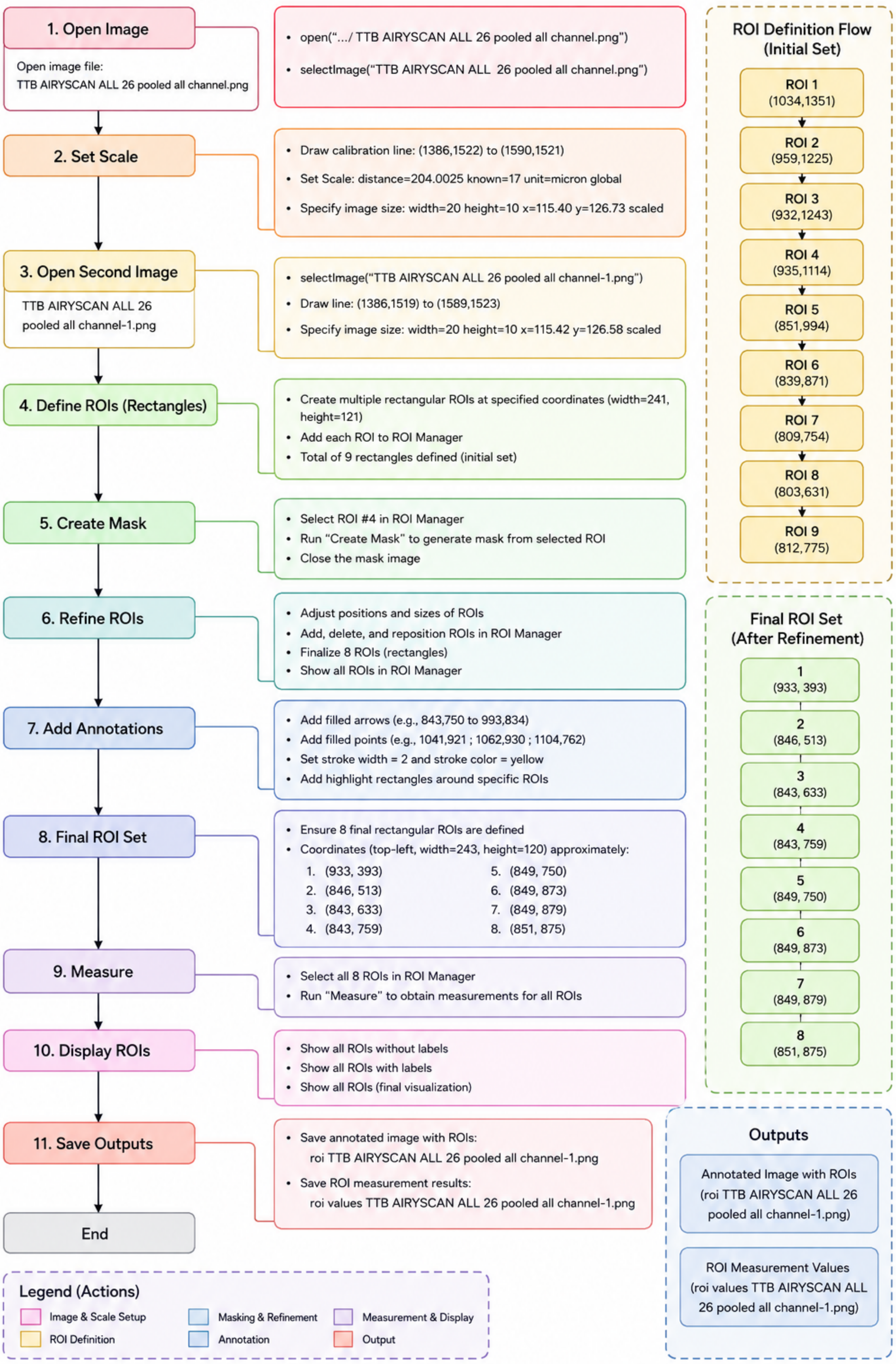
1. Open Image
Open image file:
TTB AIRYSCAN ALL 26 pooled all channel.png
open(".../ TTB AIRYSCAN ALL 26 pooled all channel.png")
selectImage("TTB AIRYSCAN ALL 26 pooled all channel.png")
2. Set Scale
Draw calibration line: (1386,1522) to (1590,1521)
Set Scale: distance=204.0025 known=17 unit=micron global
Specify image size: width=20 height=10 x=115.40 y=126.73 scaled
3. Open Second Image
TTB AIRYSCAN ALL 26 pooled all channel-1.png
selectImage("TTB AIRYSCAN ALL 26 pooled all channel-1.png")
Draw line: (1386,1519) to (1589,1523)
Specify image size: width=20 height=10 x=115.42 y=126.58 scaled
4. Define ROIs (Rectangles)
Create multiple rectangular ROIs at specified coordinates (width=241, height=121)
Add each ROI to ROI Manager
Total of 9 rectangles defined (initial set)
5. Create Mask
Select ROI #4 in ROI Manager
Run "Create Mask" to generate mask from selected ROI
Close the mask image
6. Refine ROIs
Adjust positions and sizes of ROIs
Add, delete, and reposition ROIs in ROI Manager
Finalize 8 ROIs (rectangles)
Show all ROIs in ROI Manager
7. Add Annotations
Add filled arrows (e.g., 843,750 to 993,834)
Add filled points (e.g., 1041,921 ; 1062,930 ; 1104,762)
Set stroke width = 2 and stroke color = yellow
Add highlight rectangles around specific ROIs
8. Final ROI Set
Ensure 8 final rectangular ROIs are defined
Coordinates (top-left, width=243, height=120) approximately:
1. (933, 393)
2. (846, 513)
3. (843, 633)
4. (843, 759)
5. (849, 750)
6. (849, 873)
7. (849, 879)
8. (851, 875)
9. Measure
Select all 8 ROIs in ROI Manager
Run "Measure" to obtain measurements for all ROIs
10. Display ROIs
Show all ROIs without labels
Show all ROIs with labels
Show all ROIs (final visualization)
11. Save Outputs
Save annotated image with ROIs:
roi TTB AIRYSCAN ALL 26 pooled all channel-1.png
Save ROI measurement results:
roi values TTB AIRYSCAN ALL 26 pooled all channel-1.png
End
ROI Definition Flow (Initial Set)
ROI 1 (1034,1351)
ROI 2 (959,1225)
ROI 3 (932,1243)
ROI 4 (935,1114)
ROI 5 (851,994)
ROI 6 (839,871)
ROI 7 (809,754)
ROI 8 (803,631)
ROI 9 (812,775)
Final ROI Set (After Refinement)
1 (933, 393)
2 (846, 513)
3 (843, 633)
4 (843, 759)
5 (849, 750)
6 (849, 873)
7 (849, 879)
8 (851, 875)
Outputs
Annotated Image with ROIs (roi TTB AIRYSCAN ALL 26 pooled all channel-1.png)
ROI Measurement Values (roi values TTB AIRYSCAN ALL 26 pooled all channel-1.png)
Legend (Actions)
Image & Scale Setup
Masking & Refinement
Measurement & Display
ROI Definition
Annotation
Output

86

**Semi-automated Fiji workflow for quantitative analysis of test-tube brush (TTB) assemblages.** Custom Fiji (ImageJ) macros developed specifically for this study were used to standardize the quantitative analysis of TTB architectures across confocal image datasets. The workflow begins with image loading and spatial calibration, followed by automated definition of rectangular regions of interest (ROIs), mask generation, and iterative ROI refinement. User-guided annotation was subsequently applied to identify structural features of interest before final ROI selection and quantitative measurement. The macros generated annotated images together with ROI-based measurement outputs for downstream statistical analysis. Automated processing ensured consistent application of image calibration, ROI handling, and measurement parameters across all samples while allowing minimal user intervention for quality control and refinement.
