## Supplementary material for "The Spatial Architecture of Human Subgingival Biofilms": SupplementaryInformation_SubP.pdf

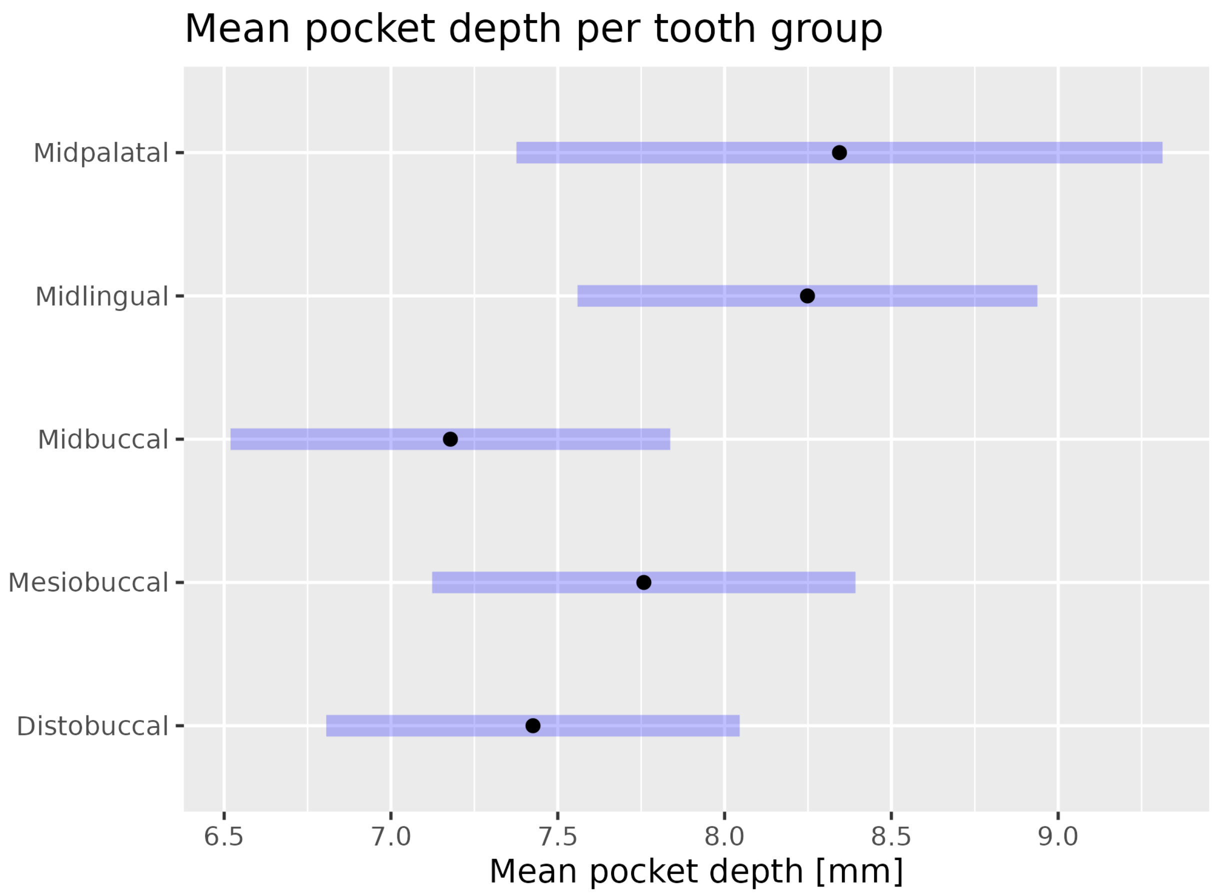

**Supplementary Fig. S1 | Model-estimated mean periodontal pocket depth across anatomical tooth groups.** Estimated marginal means of periodontal pocket depth were derived from a linear mixed-effects regression model accounting for repeated measurements within patients. Black points represent the model-estimated mean pocket depth for each anatomical tooth group, and shaded horizontal bars indicate the corresponding 95% confidence intervals. Anatomical tooth group was significantly associated with pocket depth ($\chi2=17.23$, $df = 4$, $p = 0.0017$). Midlingual sites had significantly greater estimated pocket depths than distobuccal sites, with an adjusted mean difference of 0.82 mm (95% CI, 0.04–1.60 mm; adjusted $p=0.032$), and midbuccal sites, with an adjusted mean difference of 1.07 mm (95% CI, 0.24–1.90 mm; adjusted $p=0.004$). No other pairwise comparisons were statistically significant after adjustment for multiple comparisons.

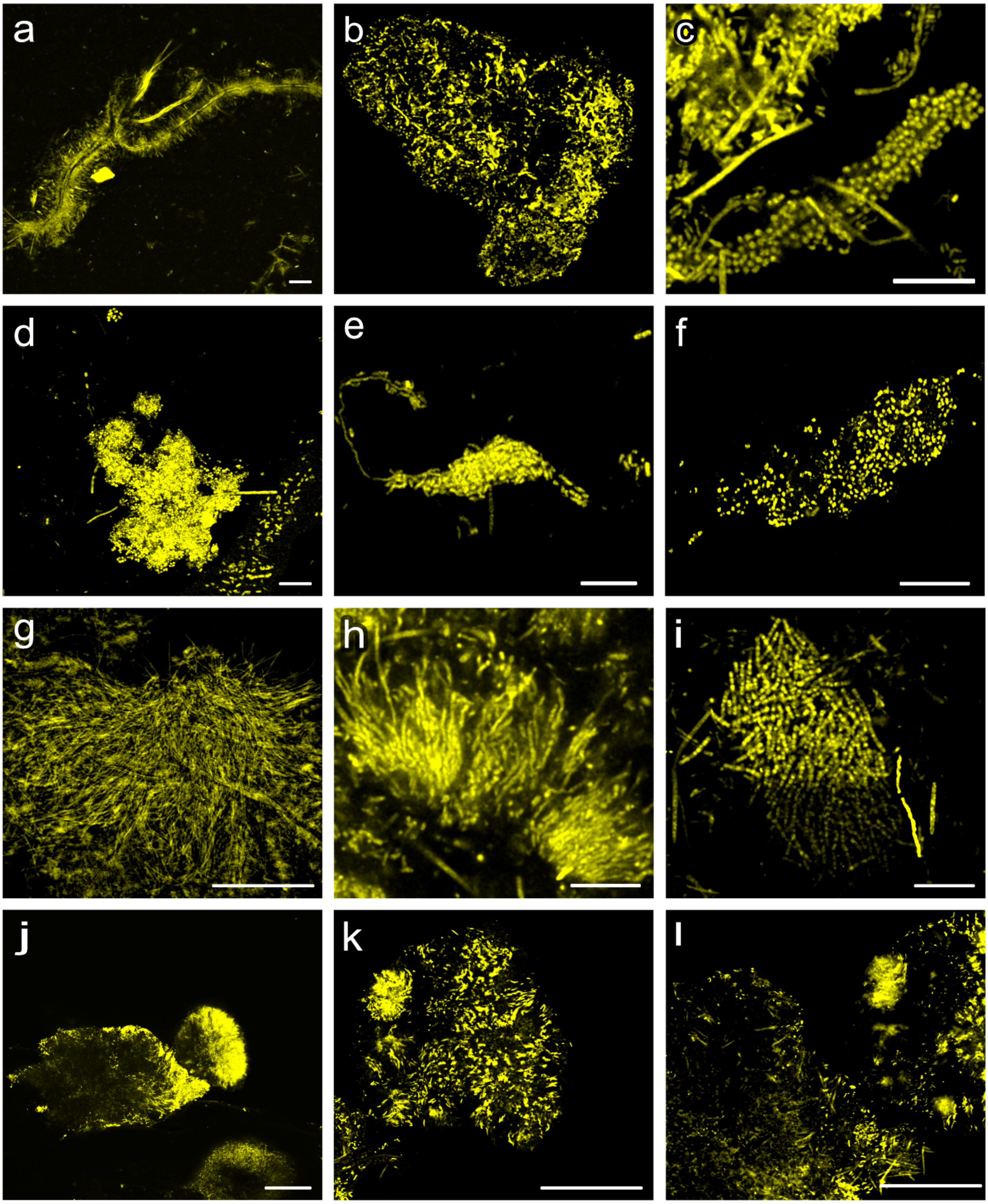

**Supplementary Fig. S2 | CLSM images of SubP stained with a universal bacterial FISH probe reveal a variety of spatially organized microbial structures across patient samples.** Panels show **a)** TTB structure, **b)** *Pg-Fn-Td* cluster, **c)** corncob**, d)** *Achillea*-like cluster, **e)** dragon-like assemblage, **f)** *Prevotella* and *Fa* dense microcolony, **g)** streaming biofilm (bio-stream**), h)** *echidna*-like assemblage, **i)** *Act*-cluster, **j-l)** amorphous orbs. Scale bars: 10 µm.

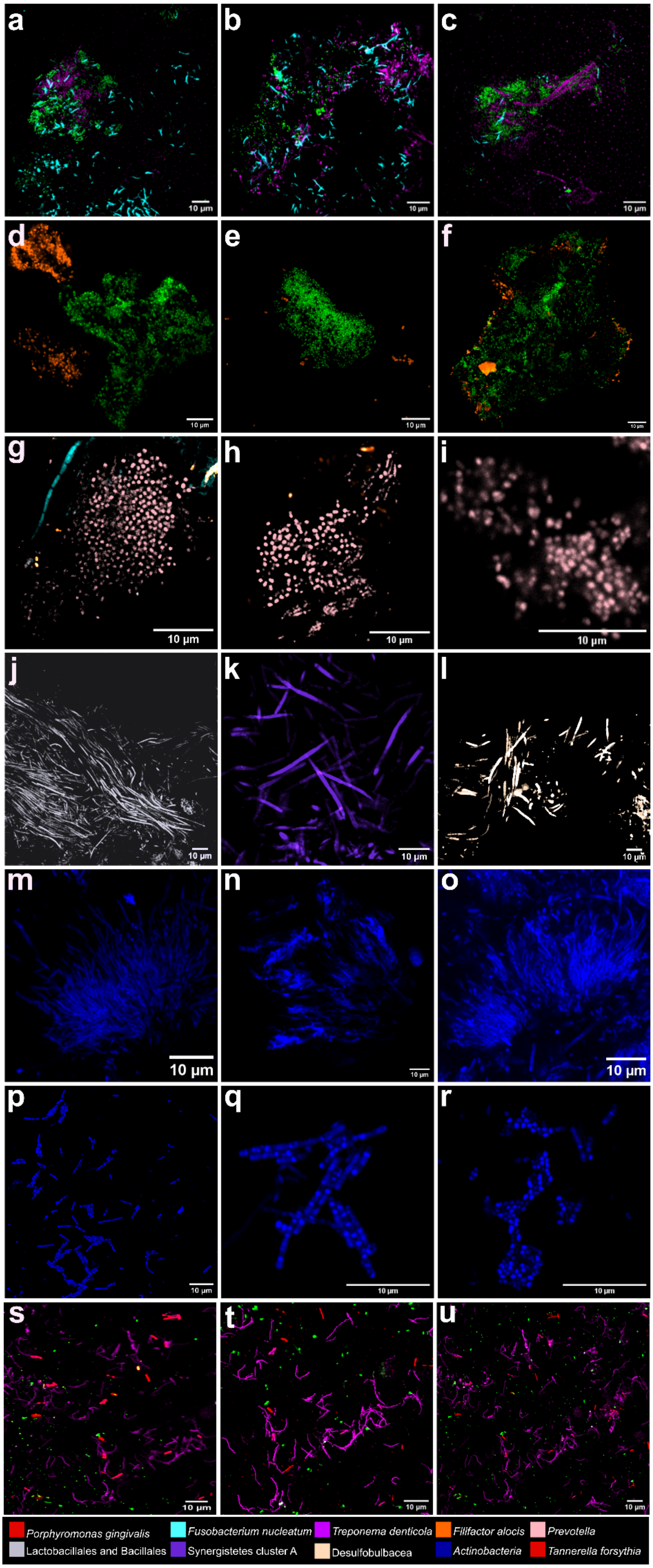
a
b
c
d
e
f
g
h
i
j
k
l
m
n
o
p
q
r
s
t
u
10 µm
Porphyromonas gingivalis
Fusobacterium nucleatum
Treponema denticola
Filifactor alocis
Prevotella
Lactobacillales and Bacillales
Synergistetes cluster A
Desulfobulbacea
Actinobacteria
Tannerella forsythia

**Supplementary Fig. S3 | Morphological and taxonomic diversity of subgingival plaque assemblages revealed by multiplexed FISH.** Panel shows CLSM images of SubP stained with taxon-specific FISH probes. **a–c,)** *Pg-Td-Fn* clusters in which *Pg, Td and Fn* co-localize within the same structure. **d–e**) *Achillea*-like clusters characterized by a dense *Pg* core surrounded by peripheral *Fa*. **g–i),** Compact *Prevotella* microcolonies, sometimes associated with one or two *Fa* cells. **j–l,** Biostreams predominantly formed by Lactobacillales and Bacillales (**j**), Synergistetes cluster A (**k**), and rarely by Desulfobulbaceae (**l**). **m–o**) *Echidna*-like assemblages composed of Actinobacteria with radial filamentous extensions, **p–r**) *Act*-clusters formed by Actinobacteria, displaying variation in cell shape from coccoid to elongated morphologies, **s–u)** Co-localization of the red complex—*Pg*, *Td*, and *Tf*—was observed across multiple fields in several patients. These taxa were abundant but typically arranged as dispersed associations rather than forming thick biofilms or dense structures. They were most frequently detected at 9 mm sites in non-smokers and at 10–12 mm sites, including 34 Lingual, 26 Mesial, 27 Lingual, 16 Distal, and 45 Mesial. Scale bars: 10 μm.

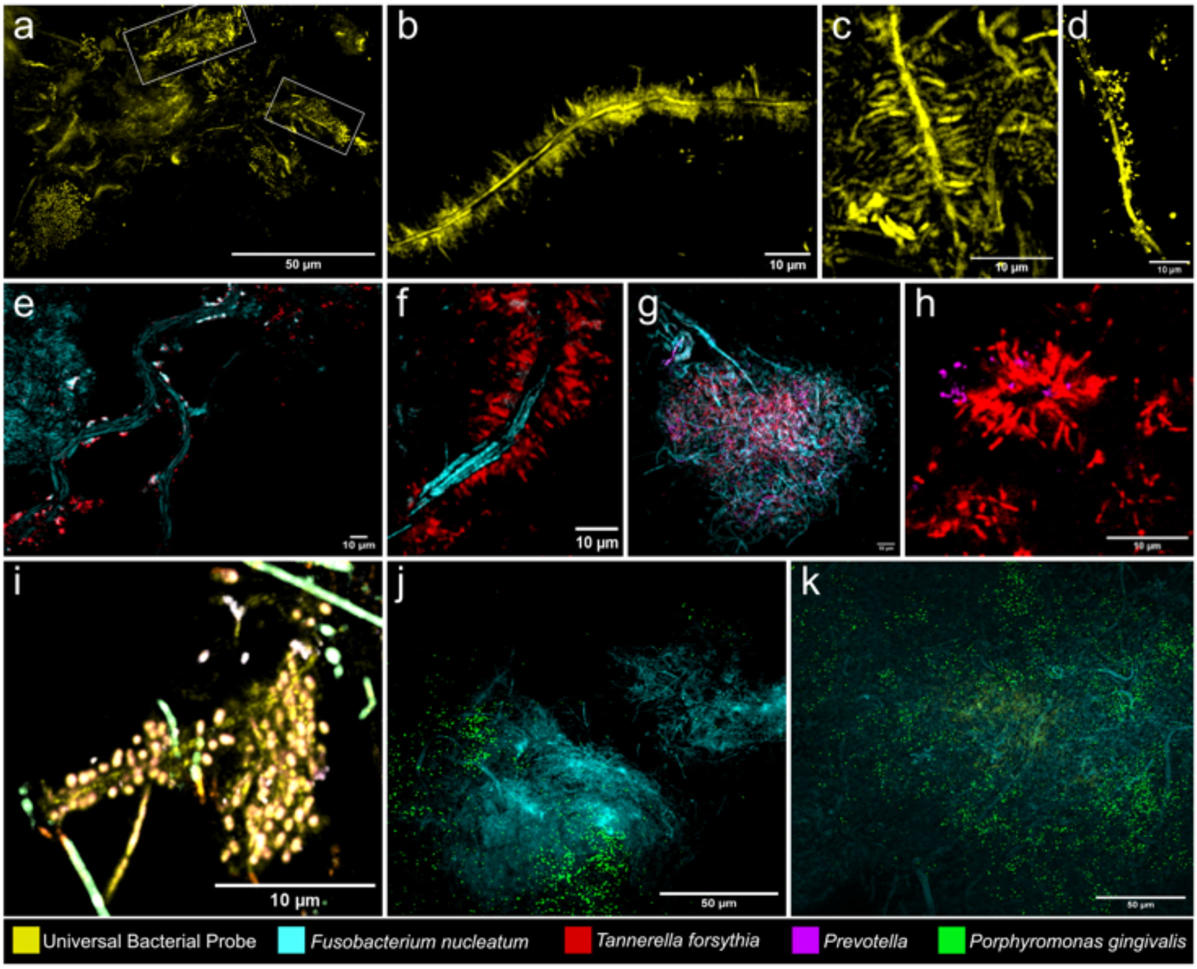

**Supplementary Fig. S4. | Spatial diversity and species composition of TTB structures in SubP from deep periodontal pockets (9-13mm). a–d)** Single-channel mFISH with the universal bacterial probe (yellow) highlights the overall TTB architecture, showing central filaments (CFs) 20–150 µm in length with orthogonally arranged bristle-like cells. **e,f)** Species-resolved imaging identifies *Fn* (cyan) as a frequent CF constituent with *Tf* (red) forming the bristles. **g,h**) Bristles are occasionally co-colonized by *Prevotella* spp. (magenta). (**i**) Representative TTB with a CF not formed by *Fn* (appears yellow; eubacterial probe), with bristles predominantly colonized by *Prevotella* and surrounded by Synergistetes cluster A (light green). **j,k**) *Pg* (green) appears in different planes of Z-stacks but is rarely closely integrated with *Tf*–*Fn* core. Scale bars: 50 µm (a,j,k), 10 µm (b–i).

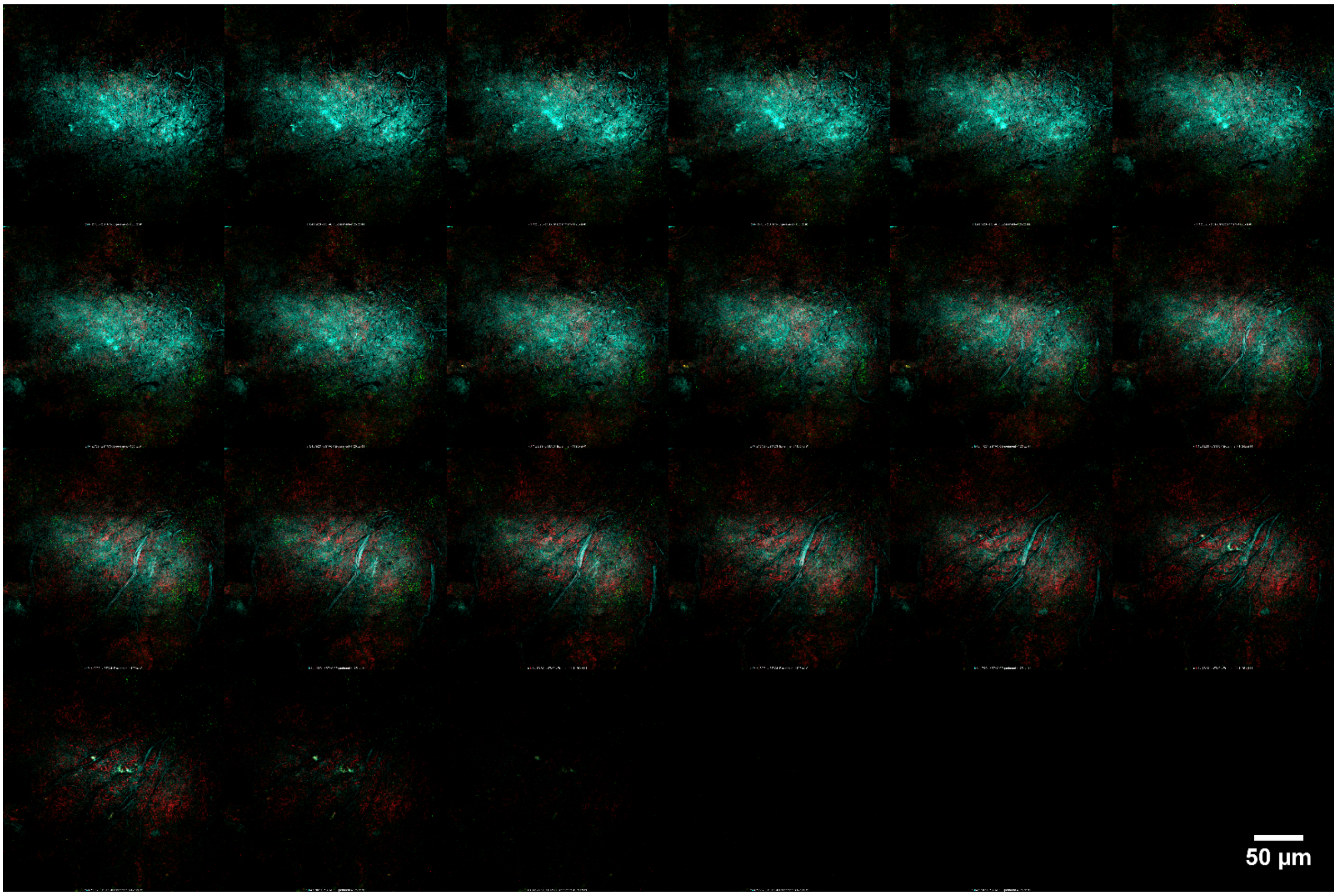

**Supplementary Fig. S5. |** Montage of the TTB Z-stack shown in supplementary Movie 2. A montage of the 22-slice (~9 µm) Z-stack highlights the species level composition of the TTB. *Fn* (cyan), *Tf* (red) and *Pg* (green).

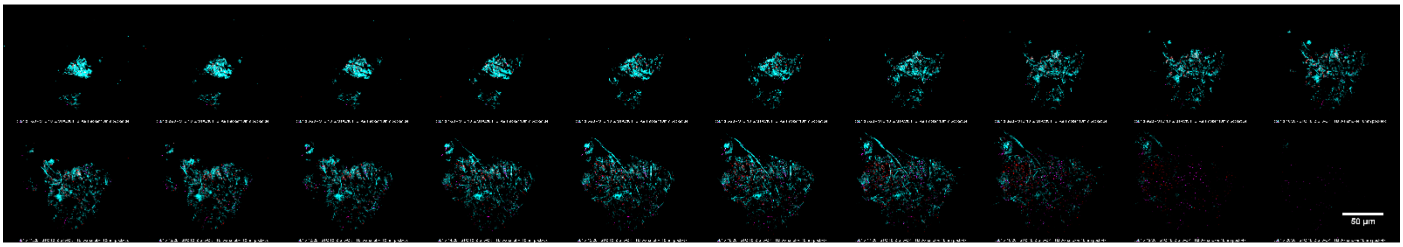

**Supplementary Fig. S6. |** Montage of the TTB Z-stack shown in supplementary Movie 3. A montage of the 20-slice (~8 µm) Z-stack highlights the species level composition of the TTB. *Fn* (cyan), *Tf* (red) and *Prevotella* (magenta). Scale bar: 50 µm.

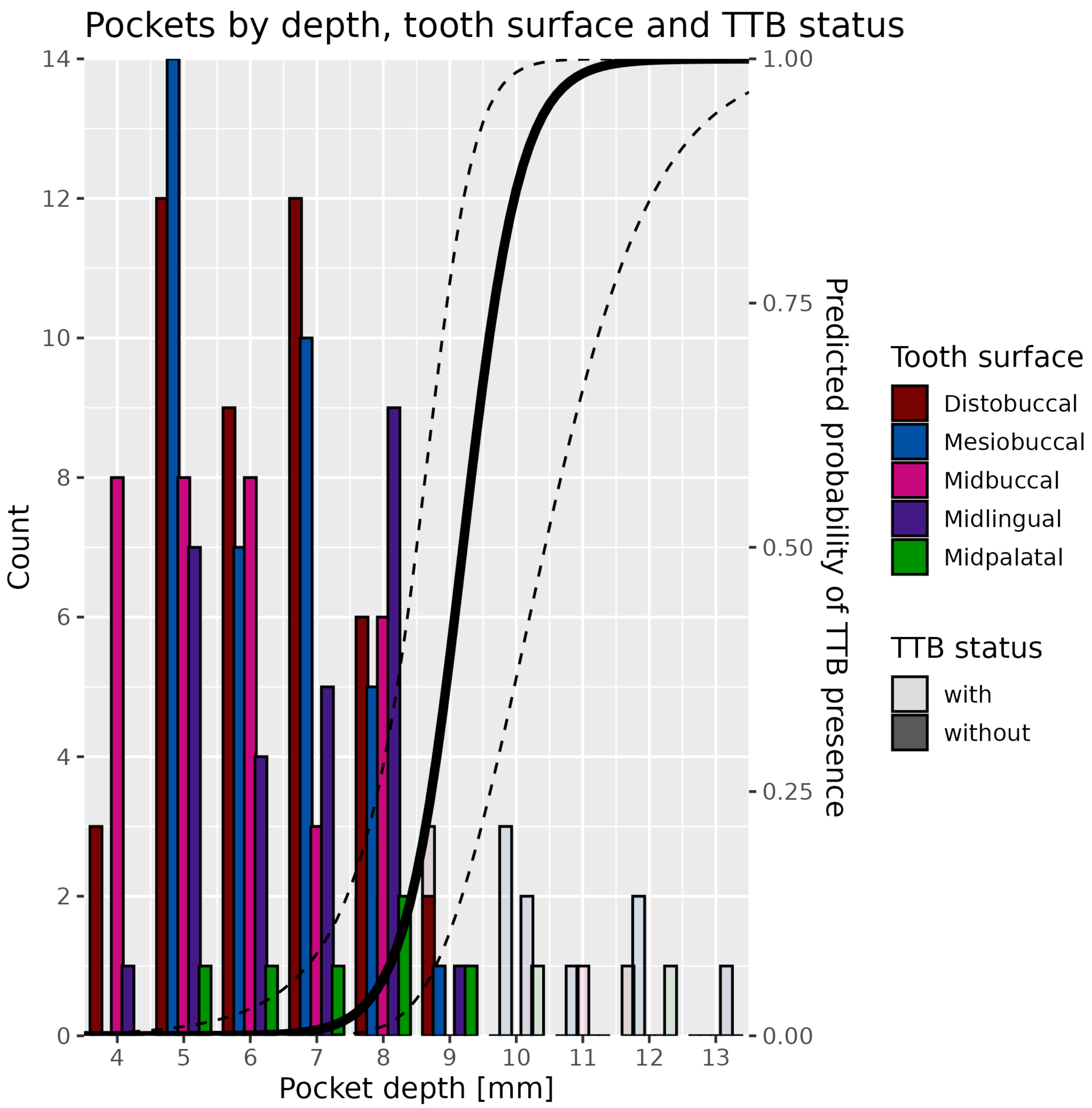

**Supplementary Figure S7 | Distribution of TTB-positive and TTB-negative sites across pocket depth and tooth surface.** Histogram showing the number of sampled subgingival sites stratified by recorded pocket depth, anatomical tooth surface, and TTB detection status. Bars represent site counts at each pocket depth and are coloured by tooth surface. Bar shading indicates whether TTB structures were detected at the site (light shade- with TTB and dark shade- without TTB). The fitted logistic curve shows the modelled relationship between pocket depth and probability of TTB detection, with dashed lines indicating the 95% confidence interval. TTB-positive sites were enriched in deeper periodontal pockets, supporting the association between increasing pocket depth and site-level TTB occurrence.

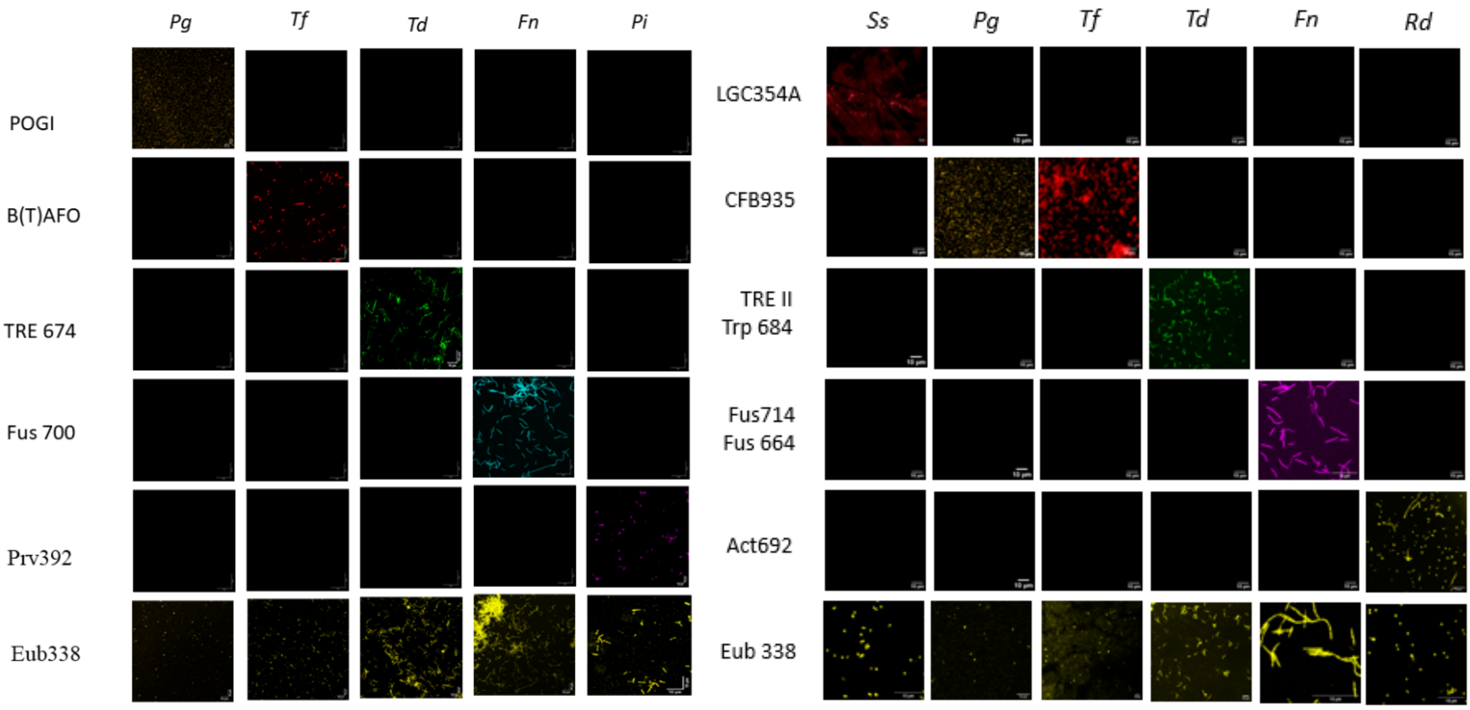

**Supplementary Figure S8 |** The effectiveness and specificity of each probe were evaluated through hybridization experiments using pure cultures of both target and non-target taxa. Simultaneously, each culture was subjected to hybridization with a universal bacterial probe for comparison. The newly designed probes were confirmed to be specific as the expected probe signals were observed for the target taxa. Scale bars- 10µm.

### Supplementary Tables

**Supplementary Table 1 (attached separately): Clinical and site metadata for subgingival plaque (SubP) samples.** This table summarises the clinical and demographic metadata associated with each subgingival plaque sample included in the study. Recorded variables include patient ID, bleeding on probing status, periodontal pocket depth, tooth position and sampling site, age, gender, relevant clinical history, smoking status, and diabetes mellitus status. Each row represents an individual sampling site.

**Supplementary Table 2**. **Bacterial assemblages in SubP with associated site, depth, smoking status, and frequency.** DB, distobuccal; MB, mesiobuccal; MIB, midbuccal; MIL, midlingual. Please refer to extended data figure 1a for the site annotation details.

| **Assemblage Type** | **Figure** | **Tooth-Site(s)** | **Pocket Depth(s) (mm)** | **Smoking Status** | **Frequency** |
|---|---|---|---|---|---|
| *Pg-Td-Fn* cluster | 1e | 13DB, 12DB, 15MIL, 17MIL, 16DB | 7, 9, 12 | Non-smokers; light & heavy smokers | Multiple |
| Corncob-like assemblage | 1f | 23MB | 10 | Heavy smokers | One FOV in 174 sites |
| *Achillea*-like floral clusters | 1g | 26DB, 17DB, 11MB, 21MB, 26MIB, 14DB, 35DB | 5–9 | Non-smokers; light smokers (only in 5 mm) | Multiple |
| Dragon-like assemblage | 1h | 26MB | 12 | Heavy smokers | One FOV in 174 sites |
| *Prevotella* microcolonies | 1i | 45MIL, 24MIL, 36DB | 9 | Light & heavy smokers | Multiple |
| Biostreams | 1j | 14DB, 16DB, 45MIL, 26MIL | 6–12 | Non-smokers; light & heavy smokers | Multiple |
| *Echidna*-like assemblage | 1k | 26MB 27DB 16MB 26MIL | 7–9 | Light smokers | Multiple |
| *Act*-clusters | 1l | 26MIL, 21MIB | 7–9 | Non-smokers | Multiple |

| Parameters of TTBs<br>Sl. No. | Total Length of TTB | Type and Length of CFs | Composition of CFs | Composition of Bristles |
|---|---|---|---|---|
| 1 | Macro TTB<br>(~ 60-150 µm) | Single long<br>(~60-100 µm)<br>(Fig. 2a; Supplementary Fig. S4 a) | Unidentified | Unidentified |
| | | Multiple short<br>(20-30 µm)<br>(Fig. 2b; Supplementary Fig. S4b, e) | *Fusobacterium nucleatum* | *Tannerella forsythia* and other taxa |
| 2 | Meso TTB<br>(40-60 µm) | Single long<br>(40-60 µm)<br>(Fig. 2c) | *Fusobacterium nucleatum* | *Tannerella forsythia* and other taxa |
| | | | Unidentified | *Tannerella forsythia* and *Prevotella* |
| | | Multiple short<br>(20-30 µm)<br>(Fig. 2f; Supplementary Fig. S4.f) | Unidentified | Unidentified |
| | | | *Fusobacterium nucleatum* | *Tannerella forsythia* and other taxa |
| 3 | Micro TTB<br>(below 40 µm) | Single long<br>(20-40 µm)<br>(Figure 2g; Supplementary Fig. S4c, d,i) | Unidentified | *Prevotella sp with* Synergistis cluster A |

**Supplementary Table 3. Classification of Test Tube Brush (TTB) Structures Based on Total Length, Central Filament Morphology, and Bacterial Composition**

**Supplementary Table 4: Summary of TTB structure counts, average lengths, total cumulative lengths, and percentage contributions based on observations across clinical samples**

| TTB Type | Number of TTBs | % of Total TTBs | Average Length per TTB (μm) | Total Length (μm) | % of Total Length |
|---|---|---|---|---|---|
| Macro-Single Long | 10 | 13.5% | 90 | 900 | 26.3% |
| Macro-Multiple Short | 6 | 8.1% | 25 | 150 | 4.4% |
| Meso-Single Long | 38 | 51.4% | 50 | 1900 | 55.6% |
| Meso-Multiple Short | 14 | 18.9% | 25 | 350 | 10.2% |
| Micro-Single Long | 6 | 8.1% | 20 | 120 | 3.5% |

A total of 74 TTBs were identified and classified into five structural categories: Macro-Single Long, Macro-Multiple Short, Meso-Single Long, Meso-Multiple Short, and Micro-Single Long. The number of TTBs per category was recorded, and the representative average filament length for each subtype was estimated from measurements of representative images: 90 μm for Macro-Single Long, 25 μm for Macro-Multiple Short, 50 μm for Meso-Single Long, 25 μm for Meso-Multiple Short, and 20 μm for Micro-Single Long. The estimated cumulative length for each category was calculated by multiplying the number of structures by the representative average length assigned to that subtype. To characterize the relative contribution of each TTB subtype, percentages were calculated relative to both the total number of classified TTBs and the total estimated cumulative TTB length across all categories.

**Supplementary Table 6. Detailed list of target taxa and probes with the corresponding fluorophores that are used for the screening of the patient derived SubP samples.**

| Sl. No | Taxon | Main Target Taxa in SubP | Probe name | Sequence (5’-3’) | Reference |
|---|---|---|---|---|---|
| | **Phylum** | | | | |
| 1. | Phylum Actinobacteria | *Rothia*<br>*Corynebacterium*<br>*Actinomycetes* | Act692 | CTGATATCTGCGCATTCC | [1] |
| 2. | Cytophaga-Flavobacterium-Bacteroides cluster | *Phorphyromonas*<br>*Tanerealla*<br>*Prevotella*<br>*Flavobacterium*<br>*Capnocytophaga*<br>*Cytophaga*<br>*Bacterioides* | CFB935 | CCACATGTTCCTCCGCTTGT | [2] |
| | **Class** | | | | |
| 3. | Class: Gammaproteobacteria | *Aggregatibacter*<br>*Morexella*<br>*Acinetobacter* | GAM42a | GCC TTC CCA CAT CGT TT | [3] |
| | **Order** | | | | |
| 4. | Lactobacillales and Bacillales | *Streptococcus*<br>*Granulicatella*<br>*Gemmella*<br>*Lactobacillus*<br>*Staphylococcus* | LGC354A | TGGAAGATTCCCTACTGC | [4] |

| | | | | | |
|---|---|---|---|---|---|
| | **Family** | | | | |
| 5. | Veillonellaceae | *Anaeroglobus*<br>*Veillonella*<br>*Megasphaera* | Prop853 | ATTGCGTTAACTCCGGCAC | [5] |
| 6. | Lachnospiraceae | *Johnsonella*<br>*Catonella* | LAC435 | TCTTCCCTGCTGATAGA | [6] |
| | **Genus** | | | | |
| 7. | | *Fusobacterium* | Fus714 | | [7] |
| 8. | | *Treponema* | Trp684 | TCTACAGATTCCACCCCTAC | [8] |
| 9. | | *Selenomonas* | Sel60 | TCATTCGCTCCGTTCGAC | [9] |
| 10. | | *Prevotella* | Pvr394 | GCACGCTACTTGGCTGG | |
| | **Others/Species Specific** | | | | |
| 11. | Synergistetes cluster A | *Fretibacterium*<br>*Pyramidobacter* | SynA1409 | ACACCCGGCTCGGGTGGT | [2] |
| 12. | | *T. forsythia* | B(T)AFO | CGTATCTCATTTTATTCCCCTGTA | [10] |
| 13. | | *F. nucleatum* | FUS664 | CTTGTAGTTCCGC/TACCTC | [10] |
| 14. | | *P. gingivalis* | POGI | CAATACTCGTATCGCCCGTTATTC | [10] |
| 15. | | *T. denticola* | TRE III | GCTCCTTTCCTCATTTACCTTT | [10] |
| 16. | | *Filifactor alocis* | FIAL | TCTTTGTCCACTATCGTTTTGA | [11,12] |
| 17. | Domain Archaea | Most Archaea | ARC915 | GTGCTCCCCCGCCAATTCCT | [10] |
| 18. | Kingdom Fungi | Pan fungal | | CTCTGGCTTCACCCTATTC | [10] |

**Supplementary Table 7. Probe sets used for this study**

| Probe Set | Focus / Purpose | Probe Name (if known) | Fluorophore |
|---|---|---|---|
| Set 1 | Core periodontal pathogens & structure | CFB935 | Alexa Fluor (AF) 405 |
| | | POGI | AF 488 |
| | | B(T)AFO | AF 532 |
| | | Pvr394 | RRX |
| | | EUB338 | Dy 615 (dual tagged) |
| | | FUS664 | AF 647 |
| | | Fus714 | AF 700 |
| Set 2 | Secondary subgingival taxa | EUB338 | AF 488 |
| | | GAM42a | AF 555 |
| | | Prop853 | Texas Red X |
| | | LAC435 | Dy 615 |
| Set 3 | Expanded periodontal pathogens | CFB935 | AF 405 |
| | | EUB338 | AF 488 |
| | | POGI | AF 555 |
| | | Pvr392 | Rhodamine Red X |
| | | B(T)AFO | AF 647 |
| | | Trp684 | AF 700 |
| Set 4 | Alternative pathogenic targets | POGI | AF 488 |
| | | TRE III | AF 555 |
| | | Pvr394 | RRX |
| | | FIAL | AF 647 |
| | | B(T)AFO | AF 700 |
| | | EUB338 | Dy 615 |
| Set 5 | Corncob structure profiling | CFB935 | AF 405 |
| | | ACT692 | AF 488 |
| | | GAM42a | AF 555 |

| | | | |
|---|---|---|---|
| | | LGC354A | AF 647 |
| | | EUB338 | Dy 615 |
| Set 6 | TTB profiling | Act692 | DY-415 (dual tagged) |
| | | EUB338 | AF 488 |
| | | B(T)AFO | Atto 532 |
| | | Prev394 | Rhodamine Red X |
| | | Sel60 | Texas Red X |
| | | Fus714 | Alexa Fluor 647 |
| | | SynA1409 | Alexa Fluor 700 |
| Set 7 | Extracted tooth root SubP analysis | POGI | AF488 |
| | | B(T)AFO | Atto 532 |
| | | TRE III | Pacific Blue |
| | | Sel60 | Rhodamine Red X |
| | | EUB338 | Dy615 (dual tagged) |
| | | FUS664 | Alexa Fluor 700 |

**Supplementary Table 8. Summary of statistical analyses, data levels and sample sizes used in this study.** Summary of statistical tests and models used in the study, including the data level, sample size, outcome variable, predictor or grouping variable, and statistical approach for each analysis. Sample sizes differ between analyses because clinical variables were assessed at patient, site, field-of-view or structure level, and architecture-specific analyses were restricted to samples meeting imaging and classification criteria.

| Sl. No. | Analysis/question | Data level | Sample size used | Outcome variable | Predictor/grouping variable | Statistical test/model |
|---|---|---|---|---|---|---|
| 1. | Relationship between patient-level pocket depth and bleeding | Patient | 36 patients | BoP percentage | Maximum PD per patient | Spearman's correlation |
| 2. | Effect of smoking on pocket depth | Site/patient clustered | 174 sites from 36 patients | Pocket depth | Smoking status | Mixed-effects linear regression |
| 3. | Effect of smoking on BoP | Site/patient clustered | 174 sites from 36 patients | BoP status/frequency | Smoking status | Mixed-effects logistic regression |
| 4. | Effect of age and smoking on pocket depth | Patient or site-level, specify | 174 sites from 36 patients | Pocket depth | Age, smoking status | Mixed-effects linear regression |
| 5. | Anatomical surface effect on pocket depth | Site | 174 sites from 36 patients | Pocket depth | Tooth surface | Mixed-effects linear regression |
| 6. | TTB density association with pocket depth | FOV/site | 40 sites from 8 patients with 10 FOVs per probe set | Pocket depth | TTB density, age, gender; smoking excluded | Mixed-effects linear regression |

| | | | | | | |
|---|---|---|---|---|---|---|
| 7. | TTB detection across pocket depth | Site | 174 sites from 36 patients | TTB presence/absence | Pocket depth | Mixed-effects logistic regression |
| 8. | Tooth-level distribution of TTBs | Tooth/site | 40 sites from 8 patients | normalised TTB density | Tooth site | Descriptive mapping; no inferential test unless modelled |
| 9. | Structural subtype distribution of TTBs | Structure | 74 TTBs | TTB subtype and cumulative length | Size class and central filament morphology | Descriptive classification |
| 10. | Bristle-segment taxonomic composition | Segment | 486 bristle segments from 8 patients, confirm final n | Taxonomic composition per segment | Patient/site/TTB subtype | Descriptive compositional profiling |
| 11. | TTB presence by smoking status and pocket-depth category | Site | 40 sites from 8 patients. One nonsmoker and 7 smokers with TTBs | TTB-positive site proportion | Pocket-depth category and smoking status | Descriptive proportions; Fisher's exact/logistic model if powered |
| 12. | Treatment-associated change in architecture | Paired site | 9 matched sites from 3 patients; paired pre/post pocket depth values | Architecture category: TTB-positive vs corncob-like | Pre- vs post-treatment | McNemar's chi-squared test with continuity correction |

### Supplementary Movie

**Supplementary Movie 1** | Sequential Z-stack scroll of spatial organisation of a representative Test Tube Brush (TTB). Single-channel mFISH imaging with the universal bacterial probe EUB338 (yellow) shows the overall architecture of the TTB across 14 optical slices (~6 µm total thickness). Scale bar: 50 µm.

**Supplementary Movie 2** | **Sequential Z-stack showing species-resolved architecture of a TTB.** mFISH imaging reveals *Fn* (cyan) forming the central filament, with *Tf* (red) arranged as bristle-like projections. *Pg* (green) appears intermittently across planes. Scale bar: 50 µm.

**Supplementary Movie 3** | **Sequential Z-stack showing species/genus-resolved architecture of a TTB.** mFISH imaging reveals Fn (cyan) forming the central filament, with Tf (red) arranged as bristle-like projections. *Prevotella* (genus probe, magenta) closely associated with the other two species in the Z-stack. Scale bar: 50 µm.
